# Identification of methylation-sensitive human transcription factors using meSMiLE-seq

**DOI:** 10.1101/2024.11.11.619598

**Authors:** Antoni J. Gralak, Katerina Faltejskova, Ally W.H. Yang, Clemence Steiner, Julie Russeil, Nadia Grenningloh, Sachi Inukai, Mustafa Demir, Riccardo Dainese, Cooper Owen, Eugenia Pankevich, Codebook/GRECO-BIT Consortium, Timothy R. Hughes, Ivan V. Kulakovskiy, Judith F. Kribelbauer-Swietek, Guido van Mierlo, Bart Deplancke

**Affiliations:** Laboratory of Systems Biology and Genetics, Institute of Bioengineering, School of Life Sciences, Ecole Polytechnique Fédérale de Lausanne (EPFL), Lausanne, Switzerland; Swiss Institute of Bioinformatics, Lausanne, Switzerland; Institute of Organic Chemistry and Biochemistry, Czech Academy of Sciences, Prague, Czech Republic; Computer Science Institute, Faculty of Mathematics and Physics, Charles University, Prague, Czech Republic; University of Toronto, Toronto, Ontario, Canada; Institute of Protein Research, Russian Academy of Sciences, Pushchino, Russia; Vavilov Institute of General Genetics, Russian Academy of Sciences, Moscow, Russia; Department of Medical BioSciences, Radboud University Medical Center, 6500 HB Nijmegen, The Netherlands

## Abstract

Transcription factors (TFs) are key players in eukaryotic gene regulation, but the DNA binding specificity of many TFs remains unknown. Here, we assayed 284 mostly poorly characterized, putative human TFs using selective microfluidics-based ligand enrichment followed by sequencing (SMiLE-seq), revealing 72 new DNA binding motifs. To investigate whether some of the 158 TFs for which we did not find motifs preferably bind epigenetically modified DNA (i.e. methylated CG dinucleotides), we developed methylation-sensitive SMiLE-seq (meSMiLE-seq). This microfluidic assay simultaneously probes the affinity of a protein to methylated and unmethylated DNA, augmenting the capabilities of the original method to infer methylation-aware binding sites. We assayed 114 TFs with meSMiLE-seq and identified DNA-binding models for 48 proteins, including the known methylation-sensitive binding modes for POU5F1 and RFX5. For 11 TFs, binding to methylated DNA was preferred or resulted in the discovery of alternative, methylation-dependent motifs (e.g. PRDM13), while aversion towards methylated sequences was found for 13 TFs (e.g. USF3). Finally, we uncovered a potential role for ZHX2 as a putative binder of Z-DNA, a left-handed helical DNA structure which is adopted more frequently upon CpG methylation. Altogether, our study significantly expands the human TF codebook by identifying DNA binding motifs for 98 TFs, while providing a versatile platform to quantitatively assay the impact of DNA modifications on TF binding.

## Introduction

Transcription factors (TFs) are critical for gene regulation by binding their cognate binding sites (TFBS) to modulate target gene expression^1^. The interaction between TFs and DNA tends to be mediated through DNA binding domains (DBDs) that recognize distinct DNA patterns (also called ‘DNA motifs’). Despite their crucial role in gene regulation, the TFBS for approximately 400 out of around 1,600 putative human TFs remain poorly characterized^1–3^. The largest and most diverse TF family is the Cys_2_His_2_ zinc finger proteins (C2H2 ZNFs), which constitute the majority of uncharacterized TFs. A human ZNF contains on average 11 individual zinc finger domains (ZFDs), most of which are thought to be capable of binding DNA. However, not all ZFDs necessarily make DNA contacts, which complicates the inference of binding properties, resulting in nearly one third of all ∼750 predicted ZNFs still lacking clearly defined binding motifs. Additionally, recent studies have provided evidence that DNA binding is not exclusively mediated by structured DBDs and that intrinsically disordered regions (IDRs) can also alter a TF’s sequence specificity and even affinity to DNA^4–6^. Given the difficulty in structurally modeling such IDRs, this also renders the *in silico* prediction of DNA binding motifs challenging despite major recent computational advances^7,8^. The experimental identification of protein-DNA interactions therefore remains an important outstanding challenge, as it continues to be seen as the gold standard to derive DNA binding motifs.

Another convoluting factor for defining TF binding models is the epigenetic state of DNA, specifically the methylation of CG dinucleotides, which can drastically alter the binding affinity of TFs to their respective binding sites^9,10^. For instance, TF families including bHLH and bZIP TFs are frequently repelled by CG methylation^11^, while MBDs or C/EBP have an increased affinity for methylated CGs in a specific sequence context^12,13^. Various *in vitro* methods based on protein binding microarrays (PBMs) and systematic evolution of ligands by exponential enrichment (SELEX) have been developed to define the effect of DNA modification on TF binding in high-throughput, and have revealed that CG methylation can differentially modulate binding affinities even for TFs in the same TF family^11,14,15^. These methods enable profiling TF interactions with methylated DNA, but they also retain the disadvantages of the original techniques, such as the limited combinatorial space in PBM and the surplus enrichment of the strongest binding sites in SELEX. While information about methylation specificity can to some extent also be imputed from genomic assays such as chromatin immunoprecipitation sequencing (ChIP-seq), these approaches typically lack resolution (fragment sizes generally span hundreds of base pairs) and require an indirect link between TF-associated peaks with in-depth whole genome bisulfite sequencing (WGBS) from the matched cell types, as the methylation status of pulled-down, TF-bound sequences is often not available. Additionally, ChIP-seq-derived binding motifs might be cell type- or co-factor-specific and do not necessarily reflect a TF’s ability to directly bind or evade epigenetically modified DNA.

To assign binding motifs for the remaining, uncharacterized human TFs, the Codebook and GRECO-BIT initiatives joined in a collaborative attempt to determine the DNA binding sites for the remaining putative DNA-binding TFs^16^. In this effort, a total of 394 TFs including positive controls were assayed using five orthogonal experimental DNA binding assays. The latter included ChIP-seq^17^, genomic and classical high-throughput SELEX ((G)HT-SELEX)^18^, and PBMs^16^. A fifth method is presented here: selective microfluidics-based ligand enrichment followed by sequencing (SMiLE-seq), which is a high-throughput microfluidic technique for examining TF-DNA interactions that relies on the mechanically induced trapping of molecular interactions (MITOMI)^19^ concept. This comprehensive experimental strategy aimed to leverage the strengths of each of the five methods while mitigating their respective limitations, such as validating the primary motif of a TF from the often noisy but genomic binding sites acquired by ChIP-seq with the high information content motifs from HT-SELEX. Additionally, the obtained data were analyzed with multiple motif discovery tools and conservatively hand-curated^16^. The resulting motifs were then benchmarked^20^ to identify the most robust binding models.

Here, we report the SMiLE-seq-based findings of the Codebook/GRECO-BIT collaboration, having assayed 284 putative TFs and derived binding models for 98 TFs. To address the hypothesis that some of the TFs for which no DNA binding motifs were recovered may in fact bind epigenetically modified DNA instead, we developed methylation-sensitive SMiLE-seq (‘meSMiLE-seq’), an assay to simultaneously probe a TF’s affinity to unmethylated and methylated DNA. By screening 114 putative TFs using meSMiLE-seq, we identified DNA motifs that include information about (me)CG affinity for 48 TFs. In addition, we derived motifs for several TFs that exclusively associate with methylated DNA, rationalizing why they remained orphan in the canonical DNA binding assays that were employed within the Codebook Consortium. Finally, we provide evidence that certain methylation-sensitive TFs are directed to distinct binding sites in the genome in a CG methylation-dependent manner, thus validating our *in vitro* observations.

## Results

### SMiLE-seq identifies binding motifs of 98 putative human TFs

We aimed to define the DNA binding specificities of 284 putative, uncharacterized TFs (selected through manual curation and including 3 positive controls, see **Table 1** and^16^) using SMiLE-seq^21^. In this assay, DNA libraries are added to *in vitro* translated TF-GFP fusion proteins and transferred into a microfluidic device, where binding events between the TF and DNA are captured after a single enrichment step. Subjecting the TF-bound fraction to high-throughput sequencing of eluted DNA fragments then allows for computational *de novo* motif discovery (**Supplementary Figure 1a-b**)^21^. Compared to alternative approaches such as HT-SELEX, SMiLE-seq has the advantage of capturing both strong and weak interactions without compromising the sampling space such as in PBMs^22^. However, given the single round of enrichment, SMiLE-seq is sensitive to potentially skewed nucleotide distributions in the input libraries given that binding enrichment is restricted to one round (**Supplementary Figure 2a**). Additionally, while SMiLE-seq allows capturing low-affinity binding sites, the obtained data tend to be noisier than after multiple rounds of enrichment (as is done in HT-SELEX), which typically results in motifs with lower information content.

**Table 1:** Overview of TFs that were investigated in this study.

**Table 2:** Re-curated and unique TFs differing from MEX.

To mitigate the effects of potential sequence biases in the input DNA libraries on *de novo* motif discovery, we used a one-sided Fisher’s exact test to select sequencing reads containing statistically enriched *k*-mers within the SMiLE-seq data of the 284 assayed TFs (**Figure 1a**, **Methods**). The sequences containing enriched *k*-mers were then passed to the ProBound Suite, a recently developed motif discovery pipeline that proved to be powerful in predicting binding sites across a wide range of affinities^20,23^ (**Figure 1a**).

**Figure 1.**
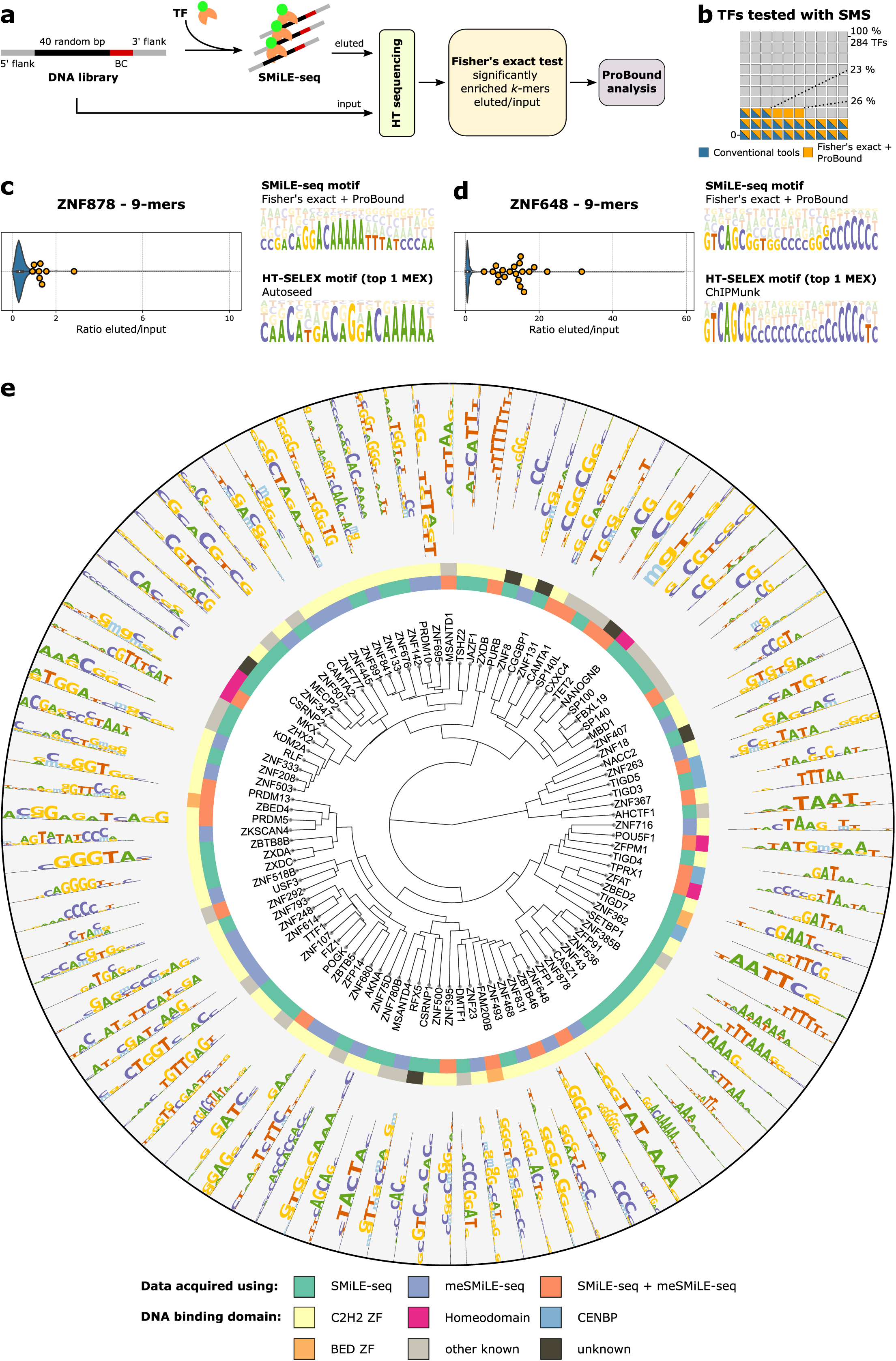
SMiLE-seq identifies DNA binding motifs for 98 putative human TFs. **a)** Schematic description of SMiLE-seq: DNA libraries are incubated with a TF of interest and transferred to the microfluidic device where interactions between DNA and TF are captured and sequenced (eluted fraction). Naïve DNA libraries are sequenced without TF enrichment (input fraction). Significantly enriched *k*-mers in the eluted fraction are identified using a one-sided, right-tailed Fisher’s exact test, with the input serving as a background distribution. Raw sequences containing significant *k*-mers are then analyzed using the ProBound Suite to infer TFBS. **b)** SMiLE-seq datasets yielded 64 DNA binding motifs when analyzed with standard motif discovery pipelines as described in^20^, whereas our analytical strategy yielded high-quality binding models for 73 TFs. **c) and d)** Violin plots depict the distributions of normalized *k*-mers in the eluted fraction (eluted/input), with the most significant *k*-mers shown as yellow dots (top 8 for ZNF878, top 20 for ZNF648). Note that these are not the most abundant *k*-mers, indicating overamplification biases in both input and eluted fractions. DNA motifs generated using the ProBound Suite after processing steps described in **a)** yield similar binding motifs as top ranked motifs reported in MEX, which were generated by orthogonal experimental methods^20^. **e)** Radial dendrogram of all reported motifs generated by SMiLE-seq and meSMiLE-seq (see below) with TF family annotation. Displayed are positive values of PSAMs. See also **Supplementary Figure 1 and 2**.

Using this approach, we successfully recovered binding motifs for 73 TFs (25.7% of all assayed TFs, including one positive control), achieving a high level of replicate reproducibility (**Supplementary Figure 2b-e**). The same data yielded only 64 approved binding models (22.5%) when analyzed without the *k*-mer enrichment step against the input library and using more conventional motif discovery tools such as Dimont, HOMER, or MEME^20,24–26^. This showcases the benefit of explicitly removing input library sequence composition bias in the data for motif discovery (**Figure 1b**). In fact, several SMiLE-seq datasets did not yield consistent motifs when analyzed with conventional tools and were thus not approved by the purposefully conservative curation and benchmarking of the Codebook Consortium. However, our analytical strategy linked 17 of these datasets to seemingly valid binding motifs, including novel models for 9 TFs that are uniquely reported in this manuscript (**Table 1** and **2**). These datasets either exhibited high replicate concordance (e.g. ZNF385B, **Supplementary Figure 2b**), or reproduced binding models generated by orthogonal experimental methods. For example, the approach extracted a similar motif for ZNF878 from SMiLE-seq data as was inferred from HT-SELEX data (**Figure 1c**), whereas straightforward processing of the same SMiLE-seq data by conventional tools failed to derive this exact motif (**Supplementary Figure 2f**). Importantly, even for some approved datasets, combining pre-filtering of reads based on *k*-mer enrichment with the ProBound Suite allowed the identification of the TF’s full-length binding site while the same datasets only yielded truncated models when analyzed with other tools, such as for ZNF648 (**Figure 1d**, **Supplementary Figure 2g** and^20^). In total, our SMiLE-seq analyses and linked analytical strategy yielded binding motifs for 50 TFs with an additional 48 derived using our methyl-sensitive SMiLE-seq method, including an overlap of 22 TFs between both approaches as detailed below. The overall 98 proteins comprise 14 different TF families, with the largest being C2H2 ZNFs (**Figure 1e**, **Supplementary Figure 2h** and **Table 1**).

**Figure 2.**
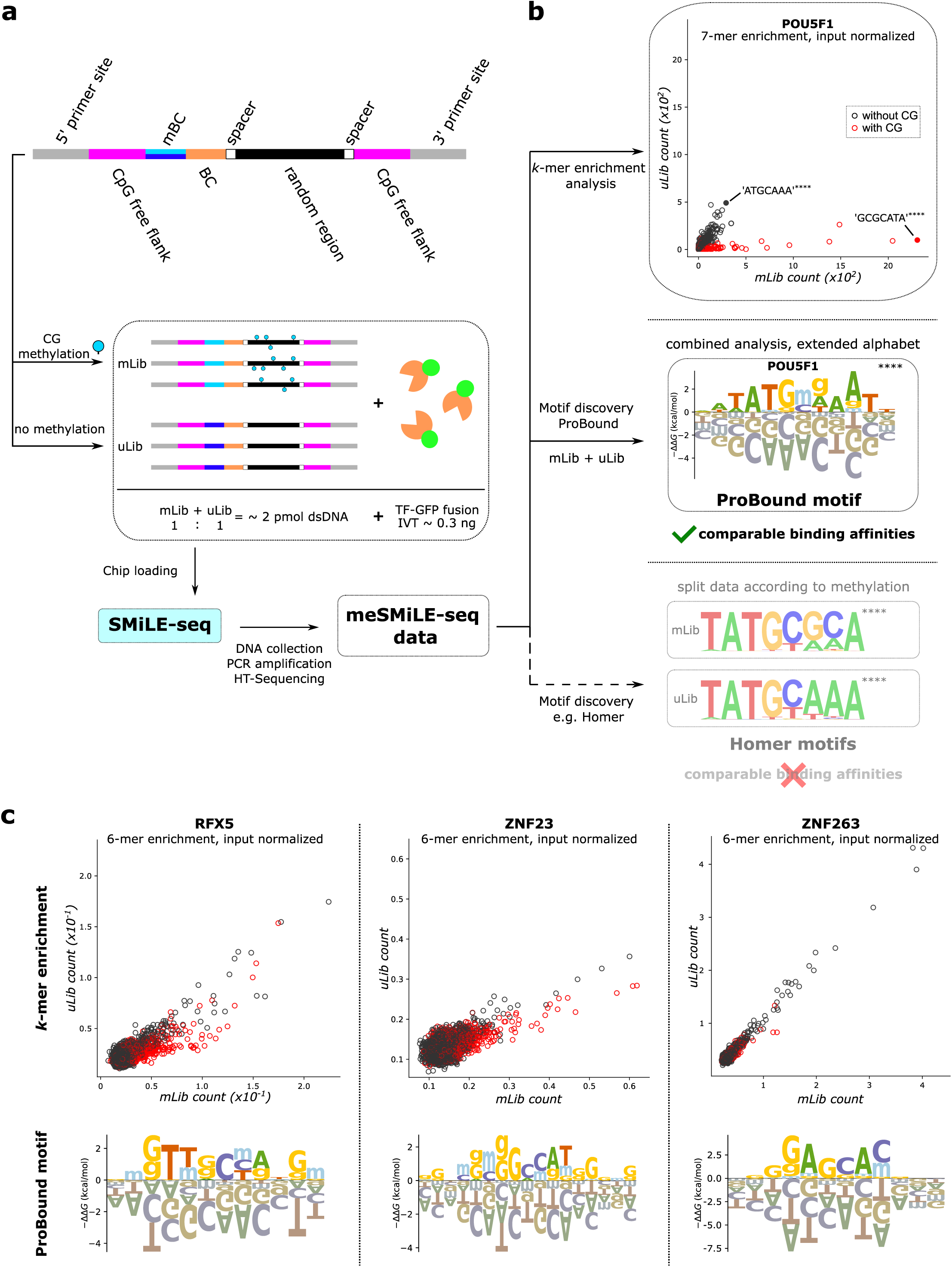
The workflow of meSMiLE-seq experiments. **a)** Each meSMiLE-seq library carries two distinct molecular barcodes (mBC and BC) and a random region comprising 24 nucleotides. To mitigate any potential affinity towards CG dinucleotides not linked to the random region, the core library is flanked by CG-free regions. According to mBC, libraries (mLib) are split and enzymatically methylated before being combined with their unmodified counterpart (uLib) and exposed to *in vitro* translated TF of interest. After a single entrapment, captured DNA is collected, amplified, and sequenced. **b) *Top panel:*** The scatterplot shows the correlation of normalized *k*-mers (eluted/input) for POU5F1 from methylated (mLib, x-axis) and unmethylated (uLib, y-axis) libraries, with each circle representing a 7-mer. Black and red colors represent the absence or presence of a CG dinucleotide within the 7-mer, respectively. Significant enrichment is tested using a one-sided Fisher’s exact test. ****P < 0.00001. ***Middle Panel:*** PSAM generated by the ProBound Suite for POU5F1. Methylation sensitivity is depicted using an extended alphabet, where ‘mg’ represents methylated CG dinucleotides. ‘g’ also indicates methylation of the complementary cytosine. ***Bottom panel:*** Two PFMs generated by HOMER for POU5F1. meSMiLE-seq data was split according to mBC. **c)** Correlation scatterplots and PSAMs (as described in **(b)**) for the three positive controls RFX5 and ZNF23 (methylation sensitive), and ZNF263 (methylation independent) as reported in Yin et al.^11^ See also **Supplementary Figure 1 and 3.**

### Development and implementation of methylation-sensitive SMiLE-seq

Despite extensive profiling attempts in our collaborative Codebook efforts, 158 putative TFs could still not be linked to binding motifs^16^. We hypothesized that some of those might prefer binding to epigenetically modified DNA instead, as the methylation of CG dinucleotides can alter a TF’s specificity and affinity for a DNA sequence^11,14^. To concurrently infer DNA binding motifs and the methylation preference of TFs, we extended SMiLE-seq to allow for a methylation-aware motif discovery (a novel workflow that we will refer to as ‘meSMiLE-seq’). Specifically, we redesigned the bait DNA libraries to expose a TF to ‘naked’ and modified DNA simultaneously (**Figure 2a**). Each DNA library thereby contains two unique molecular barcodes that provide information about the position on the microfluidic chip (BC) and its modification status (mBC). Libraries that carry a methylation barcode are enzymatically methylated prior to the experiment (methylated library, mLib) before being combined in equimolar amounts with the corresponding, unmodified counterpart (unmethylated library, uLib) and added to *in vitro* translated TF-GFP fusion proteins (**Figure 2a**, **Supplementary Figure 1**, and **Methods**). Near-complete methylation of CG dinucleotides within mLib was confirmed using digestion with the methylation-sensitive restriction enzyme *BstBI* (**Supplementary Figure 3a-b**). Considering the observed effect of input library biases in classical SMiLE-seq and the resulting challenges for data analysis, we compared different suppliers and synthesis protocols and chose input libraries with near-uniform *k*-mer distributions (**Supplementary Figure 3c-d**, **Methods**). This rigorous control removed the need to prefilter sequencing reads as was required for classical SMiLE-seq data. Since most conventional motif discovery tools were not developed to report DNA modifications, conveying position-specific information about DNA methylation using a position frequency matrix (PFM) inferred with classical tools is challenging. To overcome this issue, we used the ProBound Suite to present the predicted DNA binding motifs from the meSMiLE-seq workflow as position-specific affinity matrices (PSAM)^27^ (**Figure 2b**, **Methods**).

As a proof of concept for meSMiLE-seq, we profiled the TF POU5F1 (also known as OCT4), since it has previously been shown to bind to both methylated and unmethylated DNA motifs that direct its genomic location, making it a valuable positive control^11,28^. Using meSMiLE-seq, we found that POU5F1 enriches two distinct DNA *k*-mer species, correctly recapitulating its known genomic consensus binding sites ‘ATGCAAA’ and ‘ATG(meCG)CAT’^11,28^. The methylation-independent sequence ‘ATGCAAA’ was equally frequent in uLib and mLib, while ‘G(meCG)CATA’ was much more prevalent in mLib compared to any unmethylated sequence including ‘GCGCATA’ in uLib, showcasing the protein’s strong preference only for the methylated version of the motif (**Figure 2b**). Given meSMiLE-seq’s ‘one-pot reaction’ approach, we thereby note that our data permits a more precise estimation of actual binding preferences, as uLib and mLib are in direct competition for the TF. Therefore, *k*-mer enrichment scatterplots indicate the preferred DNA species bound by the TF. Full TFBS are shown as PSAM motifs, with letter sizes representing the stability of the TF-DNA binding complex^27^. Letters above the x-axis suggest increased stability and an extended alphabet helps to identify the impact of CG methylation on TF-DNA interactions at base pair resolution (**Figure 2b**, **Methods**). Alternatively, meSMiLE-seq data can be split according to mBC and analyzed separately using conventional approaches such as HOMER^24^, generating two independent motifs. While this strategy yields high information content motifs, it loses the possibility to directly compare binding preferences (**Figure 2b**).

Next, we assayed three more positive controls to adequately verify meSMiLE-seq. We included RFX5 and ZNF23 as controls for TFs binding to methylated sequences *in vitro*, and ZNF263 as a control for a methylation-independent binder, as all have been previously profiled for CG methylation affinity^11^. meSMiLE-seq correctly recapitulated binding sites of these TFs, enriching the methylated sequences and predicting methylation-aware motifs for RFX5 and ZNF23, and the methylation-independent motif for ZNF263 (**Figure 2c**). Together, these findings demonstrate the robustness and efficacy of meSMiLE-seq to profile methylation-dependent DNA binding specificities in parallel.

### Methylation-sensitive screening of poorly characterized human TFs

Next, we applied meSMiLE-seq to screen the DNA binding specificities of 114 poorly characterized TFs, profiling in total 128 ‘protein constructs’, i.e. 83 full-length proteins (FL) and 45 isolated DBDs (see **Table 1**, **Supplementary Figure 3e-f**). This set included 70 randomly selected Codebook protein constructs (33 FL, 37 DBDs, ‘set 1’) from approved and non-approved datasets and 23 constructs (15 FL, 8 DBDs, ‘set 2’) that previously yielded binding motifs only in ChIP-seq experiments, thus potentially implying a possible interaction with epigenetically modified DNA. In addition, we expanded this set with 35 lab-available KRAB-ZNFs (‘set 3’), since many TFs in this class display genomic binding preferences to heterochromatin-associated regions in ChIP-exo^29^, which could also suggest an ability to bind methylated DNA. Using the meSMiLE-seq pipeline, we were able to infer high-confidence binding models for 48 TFs comprising 27 TFs from ‘set 1’, 4 TFs from ‘set 2’ and 17 TFs from ‘set 3’, with detailed insights presented in the sections below (**Figure 1e**, **Supplementary Figure 3e-f**).

To confirm the validity of meSMiLE-seq-derived motifs and the robustness of the method, we used several approaches. First, we compared the meSMiLE-seq PSAMs to orthogonal data where TF specificity/affinity towards methylated DNA was not considered and we excluded the extended, methylation-specific alphabet (**Table 5**, **Methods** and **Supplementary Figure 4a**). meSMiLE-seq motifs showed high concordance with binding models predicted by orthogonal *in vitro* and *in vivo* methods, achieving an average Pearson correlation coefficient (PCC) of 0.899 across all TFs (based on values in aligned probability matrices) (**Figure 3a**, **Table 5** and^16^). We observed that TFs with longer motifs, such as PRDM10 or ZNF793, do not display strong similarity towards other TFs, even within their respective TF families (**Figure 1e**, **Figure 3a**). This was especially noticeable for C2H2 ZNF proteins, consistent with the expected uniqueness of their binding sites^29,30^. Second, we compared the meSMiLE-seq motifs of 13 C2H2 ZNFs to those derived from ChIP-seq^17,31^ and ChIP-exo data^29^ (**Table 5**) using HOMER de novo motif enrichment analysis for these same TFs (**Figure 3b**). Importantly, while most of the first-ranked (i.e. most significant reported by HOMER) ChIP-seq-inferred motifs resembled meSMiLE-seq-derived models (as indicated by the digit next to the PFMs), this was not the case for ZBTB46 (second-ranked), ZNF133 (second- and third-ranked), and ZNF445 (seventh ranked) (**Figure 3b**), illustrating the value of orthogonally validating ChIP-derived binding sites using *in vitro* assays. These findings align well with the observation that an estimated ∼25 % of all, most significant motifs reported by traditional motif discovery tools such as HOMER do not represent the ‘true’ binding sites of the studied TFs^20,32^. Instead, these motifs likely reflect those of co-factors (e.g. in ChIP-seq) or over-amplified DNA sequences due to method-related errors or biases (e.g. in HT-SELEX)^20^.

**Figure 3.**
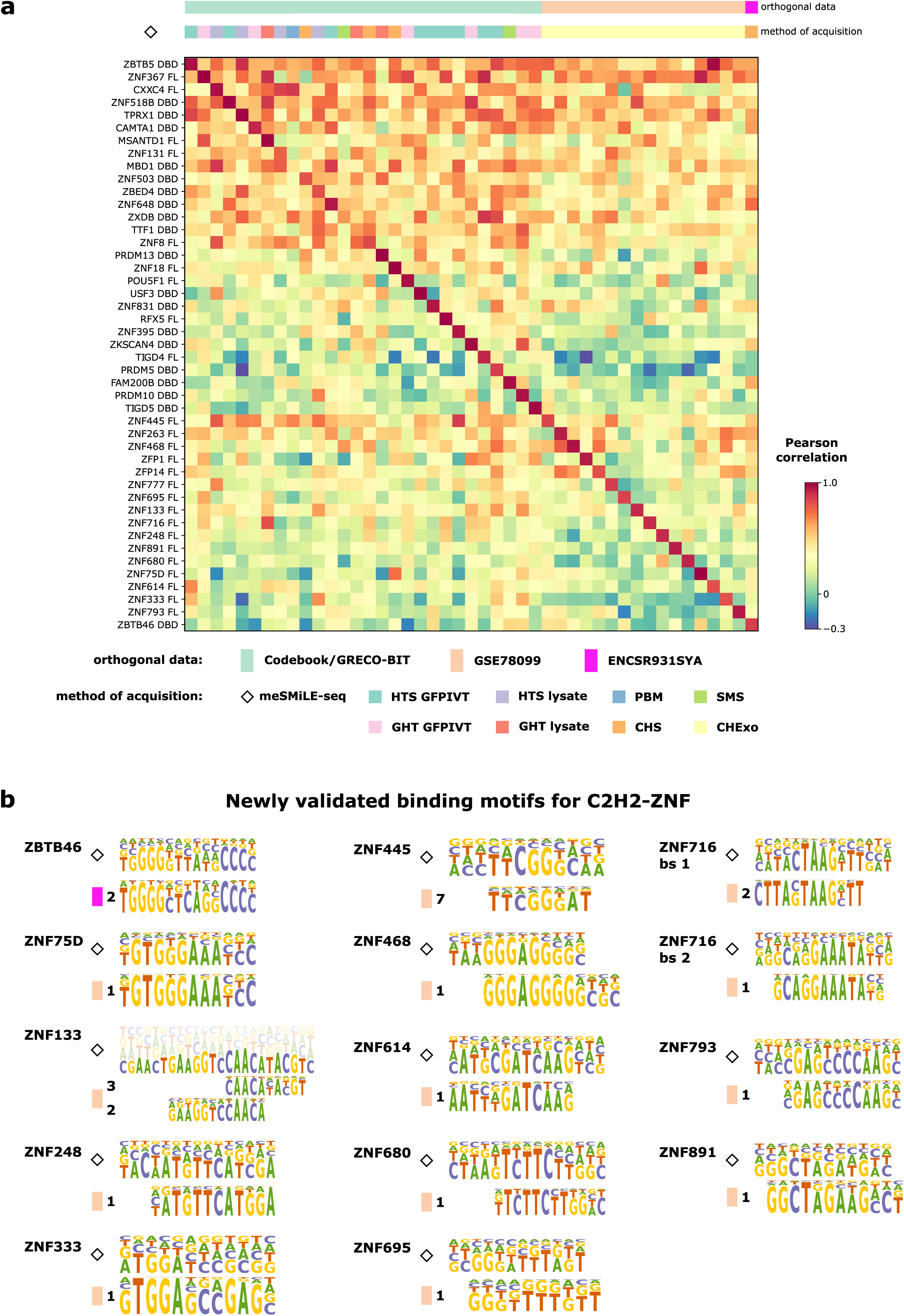
meSMiLE-seq-derived motifs correlate highly with orthogonal datasets. **a)** Correlation matrix between meSMiLE-seq-derived PFMs and DNA motifs generated by orthogonal datasets, expressed as Pearson correlation coefficients. HTS: HT-SELEX, GHT: Genomic HT-SELEX, SMS: classical SMiLE-seq, CHS: ChIP-seq, CHexo: ChIP-exo, PBM: PBM. GFPIVT: TFs expressed as GFP fusion proteins via IVT-kit, Lysate: expressed in HEK293 cells^18^ **b)** meSMiLE-seq validation of binding motifs for C2H2-ZNFs that were previously only assayed by ChIP-seq or ChIP-exo. The digit indicates the rank of the found motif generated by HOMER.

### Classification of TFs based on methylation sensitivity inferred from meSMiLE-seq

We next classified the 48 TFs with characterized binding motifs into three distinct groups depending on their affinity for methylated sequences based on ProBound predictions (see **Methods**), which we will refer to as ‘methyl plus’, ‘methyl minus’, and ‘little effect/no CG’^11^. ‘Methyl plus’ TFs (n=14) demonstrated a higher attraction towards methylated CG dinucleotides compared to unmethylated DNA, such as ZNF445, a genomic imprinter predicted to bind methylated DNA *in vivo*^33^. Additionally, some TFs from this group enriched more than one consensus sequence such as PRDM13, which binds methylation-independently to ‘GCAGGTGG’ and to methylated ‘G(meCG)GGTGG’, displaying a behavior similar to that of POU5F1. In contrast, ‘methyl minus’ TFs (n=13) had a reduced affinity to methylated DNA (e.g. USF3). TFs that interacted with sequences regardless of their methylation status or preferred motifs without CGs were classified as ‘little effect/no CG’, such as ZNF367 (**Figure 4a**, **Table 3**, **Supplementary Database 1**).

**Figure 4.**
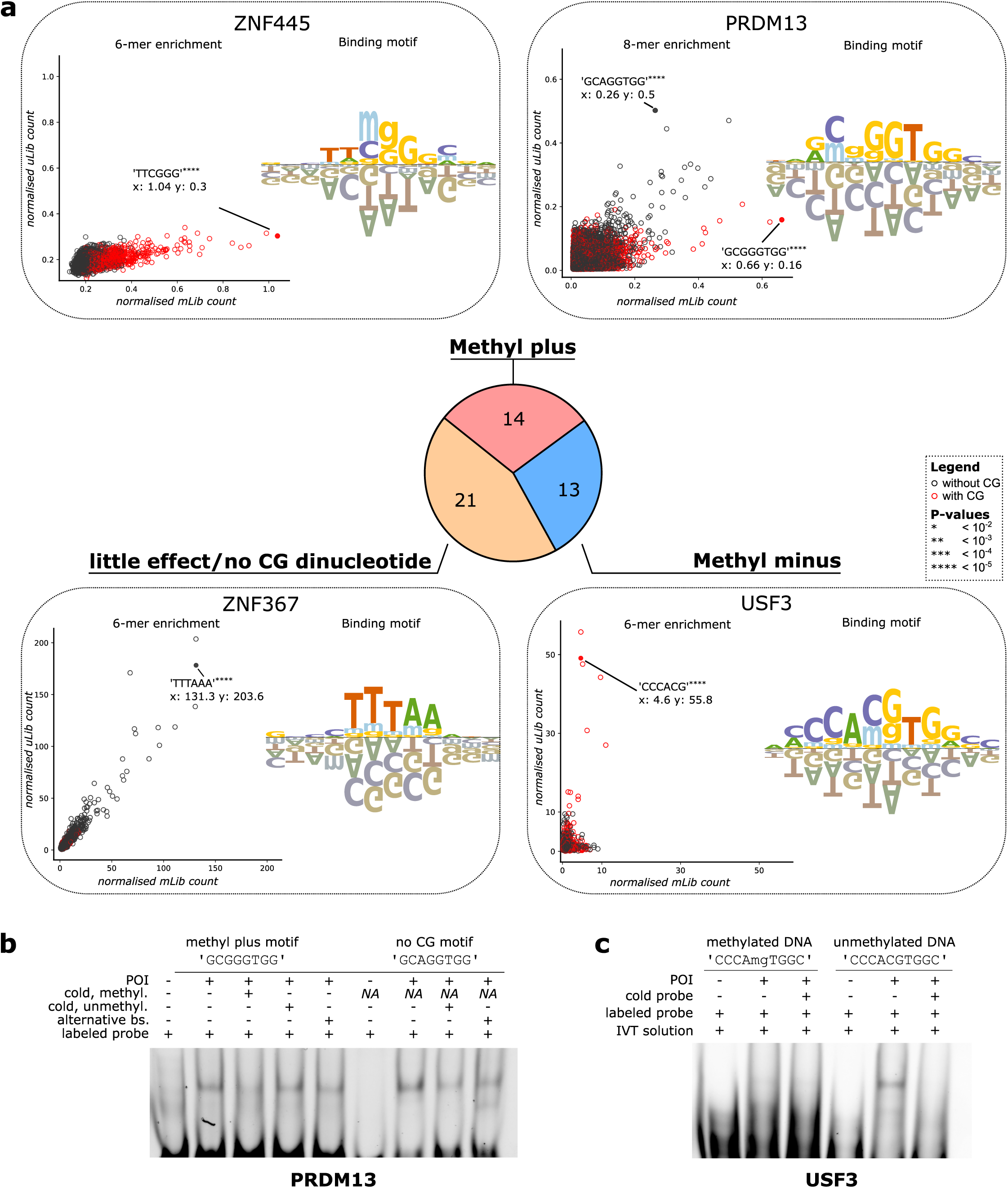
TF classification based on affinity towards methylated DNA. **a)** Categorization of TFs into three groups based on methylation sensitivity: increased affinity or alternative binding site containing methylated CG (‘methyl plus’), decreased affinity (‘methyl minus’), no observable effect (‘little effect’ or ‘noCG dinucleotide’). Depicted are correlation scatterplots and PSAMs as described in Figure 2b of exemplary TFs for each group. **b) and c)** EMSA validation of methylation-sensitive DNA binding for PRDM13 (‘methyl plus’) and USF3 (‘methyl minus’). POI: protein of interest, ‘cold, methyl.’: methylated cold probe, ‘cold, unmethyl.’: unmethylated cold probe, alternative bs.: alternative binding site (applicable for PRDM13). See also **Supplementary Figure 4b-c.**

**Table 3:** TF classification based on affinity towards methylated DNA.

To provide support for our meSMiLE-seq-derived findings, we performed electrophoretic mobility shift assays (EMSA) for PRDM13 (methyl plus) and USF3 (methyl minus). PRDM13 caused DNA shifts with both its ‘methyl plus’ and ‘no CG’ motifs, while USF3 exclusively engaged in binding with its motif when unmethylated, supporting the results from meSMiLE-seq (**Figure 4b-c**, **Supplementary Figure 4b-c**).

### The methylation of binding sites dictates the genomic distribution of TFs

Next, we investigated whether our *in vitro* ‘methylation (in)sensitivity’ assessments could be validated in a cellular context. We searched for individual occurrences of meSMiLE-seq-derived motifs for each TF within corresponding ChIP-seq^17^ or ChIP-exo^29^ data in HEK293 and HEK293T cells (**Table 5**). To obtain information on the degree of methylation at the genomic loci that contained relevant motifs, we intersected the regions with publicly available WGBS data for both cell lines^34,35^ (**Table 5**). We first focused on PRDM13, as PRDM13 binds both unmethylated and methylated DNA in different sequence contexts *in vitro* and both of these binding sites were abundantly found in ChIP-seq peaks (**Supplementary Figure 5a**). The lack of CG dinucleotides in the ‘no CG’ motif ‘GCAGGTGG’ led to the analysis of CG methylation across entire peaks containing this motif. Here, ∼45% of CGs were methylated at less than 10%, while ∼25% showed methylation levels above 90% (**Figure 5a-b**). In contrast, more than 38% of occurrences of PRDM13’s ‘methyl plus’ motif ‘GCGGGTGG’ within ChIP-seq peaks were found to be highly methylated (> 90%) in HEK293 cells. This observation is particularly striking given that CG methylation levels across entire peaks containing the ‘methyl plus’ motif are predominantly unmethylated, with ∼64% showing less than 10% methylation and only ∼14% exceeding the 90% methylation threshold (**Figure 5a-b**, **Methods**). Thus, these findings indicate that PRDM13 can bind both meSMiLE-seq predicted motifs in a methylation-dependent context *in vivo*. We expanded the analysis to include all TFs with available ChIP-seq^17^ or ChIP-exo data^29^ and observed similar patterns for ‘methyl plus’ TFs ZNF445 (**Figure 5c**), POU5F1, ZNF716, ZNF18 and ZNF518B (**Supplementary Database 2**). Interestingly, while RFX5 successfully served as a positive control for a ‘methyl plus’ TF in meSMiLE-seq, showcasing affinity to both methylated and unmethylated motifs *in vitro* (**Figure 2c**), its genomic TFBS were not methylated in HEK293 cells, consistent with previous observations in different cell lines^36^. Similar discrepancies were noticed for the ‘methyl plus’ predicted TFs ZKSCAN4, ZNF133, ZNF503, and ZNF648, which bound predominantly unmethylated regions in HEK293/T cells (**Supplementary Database 2**). These data suggest that while a TF’s ability to bind methylated DNA *in vitro* is a prerequisite for binding the modification *in vivo*, it does not guarantee that this behavior will be observable in all cell types.

**Figure 5.**
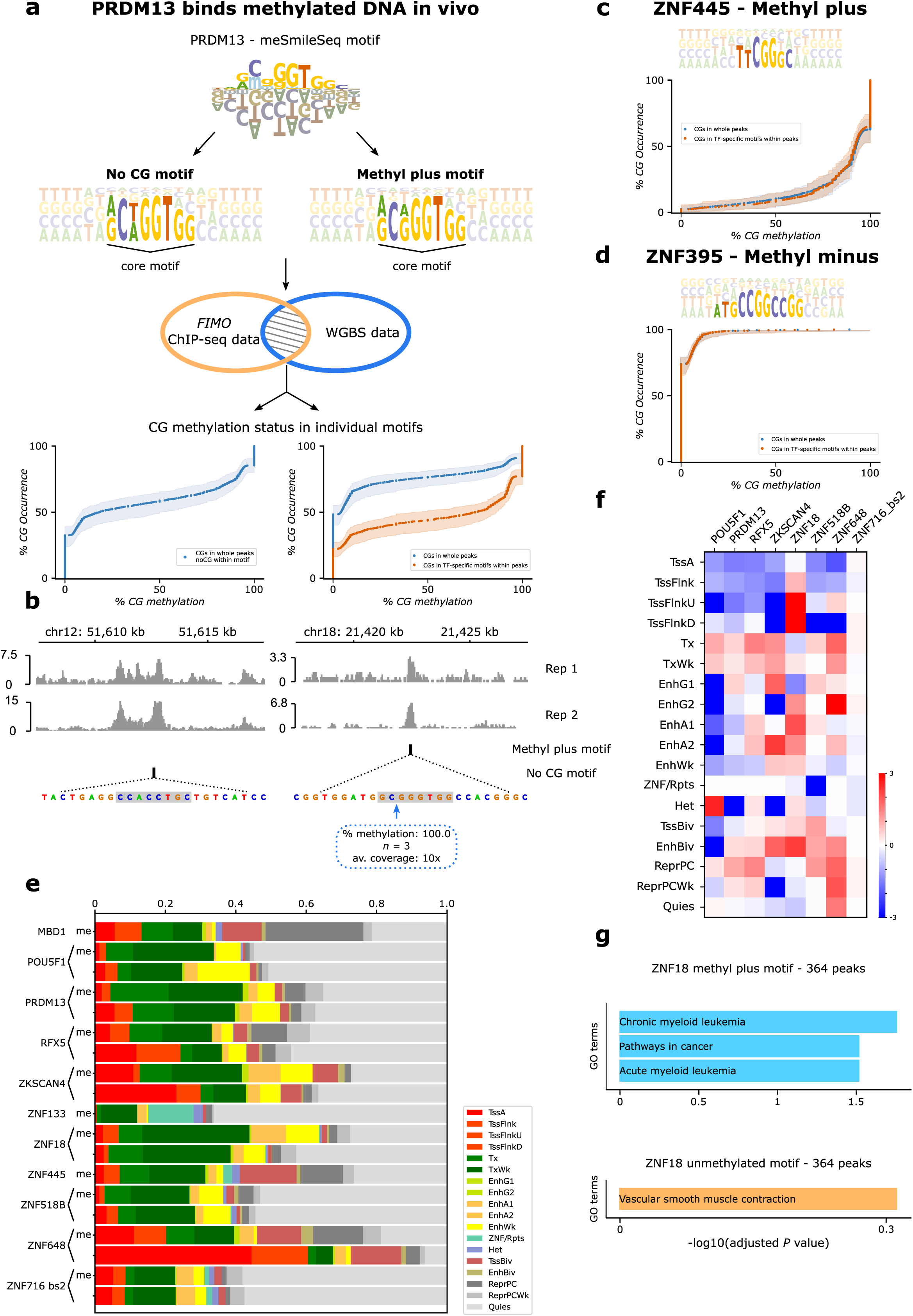
Methylation of TFBS drives genomic distribution of TFs. **a)** Exemplary workflow to identify methylated TFBS in cells. ‘methyl plus’ and ‘no CG’ motifs are extracted as PFMs from meSMiLE-seq-inferred PSAMs for PRDM13 and used to identify motif occurrences in PRMD13-specific ChIP-seq peaks. Individual instances are intersected with WGBS data, and the distribution of methylation is depicted as an ECDF. **b)** IGV browser snapshots for a ‘no CG’ and ‘methyl plus’ motif found in PRDM13-specific ChIP-seq peaks. **c) and d)** ECDFs depicting methylation levels of motifs for ZNF445 and ZNF395. **e)** ChromHMM annotations of TF-peak-specific motifs of ‘methyl plus’ TFs identified as described in (**a**). ‘me’ describes ‘methyl plus’ motifs, while the lack of ‘me’ represents ‘no CG’ or unmethylated motifs. Tss: transcription start site, TssBiv: bivalent/poised transcription start site, Tx and TxWk: actively transcribed genes, EnhBiv: bivalent enhancer. The full legend for abbreviations can be found in **Supplementary Figure 5b. f)** Heatmap of TF-specific ChromHMM annotations expressed as log2-transformed ratios (‘methyl plus’ / ‘no CG’). **g)** Gene ontology enrichment analysis of ZNF18’s ‘methyl plus’ (at least 50 % methylated) motifs (364 peaks in total) and ‘no CG’ motifs (364 most significant peaks). See also **Supplementary Figure 5a-g**.

In contrast, motif occurrences for most ‘methyl minus’ TFs (11 of 13) were predominantly unmethylated in cells, as illustrated by ZNF395 (**Figure 5d**, **Supplementary Database 2**), suggesting that these proteins might have in general a weakened affinity towards their motifs when methylated irrespective of cellular context.

Given that several TFs were found to associate with unmethylated and methylated DNA motifs with distinct sequences, we aimed to assess the genomic impact of the different binding profiles of these TFs. We intersected the binding sites of all TFs with available ChIP-seq or ChIP-exo data with ChromHMM annotations for HEK293/T cells^37^ (**Supplementary Figure 5a-c**), but focused especially on ‘methyl plus’ TFs to compare the genomic annotations of highly methylated motif occurrences (motif occurrences of ‘methyl plus’ motifs that are at least 50% methylated in WGBS) to their unmethylated counterparts (i.e. ‘noCG’ motifs) (**Figure 5e-f**). The analyses revealed that methylated binding sites are mostly depleted around active transcription start sites (Tss) (except ZNF716 and ZNF18) while being enriched in most cases in bivalent/poised regulatory elements (TssBiv and EnhBiv) and actively transcribed genes (Tx and TxWk). These observations support the methylation status of these DNA elements (**Figure 5f**) as they tend to be consistent with previous findings such as heavy methylation of gene bodies of actively transcribed genes^38–41^. Annotation of the genomic regions using gene ontology analysis of nearby genes (< 3 kb distance of the TFBS) also showed that different DNA motifs may direct TFs to distinct classes of genes and may thus contribute to the differential regulation of cellular function. This is illustrated by ZNF18, a poorly characterized KRAB-ZNF whose ZNF18 ‘methyl plus’ motifs are significantly associated with ‘pathways in cancer’ and more specifically with ‘chronic/acute myeloid leukemia’ (**Figure 5g**), which in turn might rationalize its involvement in the pathogenesis of various malignancies^42–44^. Although gene ontology enrichment for many other TFs was inconclusive as most pathways did not surpass the significance threshold (**Supplementary Figure 5d-g**), our analyses nevertheless suggest that methylation-sensitive TFs are recruited to different genomic locations based on their affinity towards DNA methylation and thus exert their regulatory function in different cellular contexts.

### ZHX2 as a putative Z-DNA binding protein

Among the detected ‘methyl plus’ TFs, we found one TF (zinc fingers and homeoboxes protein 2 (ZHX2)), that encodes two C2H2 ZNFs and four or five homeodomains (HDs)^45,46^ that specifically enriched methylated ‘CG’ repeats when expressed as a full-length protein in meSMiLE-seq. To locate the DNA interaction domains of this protein, we performed experiments using the DBDs separately and identified the HDs as mediating the observed DNA binding properties (**Figure 6a**). However, when comparing our data to that of ChIP-seq-derived ZHX2 motifs, we found that the motifs were dissimilar as the latter mainly comprised of typical promoter motifs similar to those of other promoter binders such as ‘TGACG’ or ‘AAGATGG’ for CREB1 and YY1, respectively^31,47^. Other reported, predicted genomic binding sites for ZHX2 included sequences such as ‘AGGCTAGA’^48^ or ‘CCACCAC’^49^. The methylated poly(CG) core of the ZHX2 motif derived from meSMiLE-seq is however reminiscent of DNA sequences involved in the formation of non-canonical DNA structures, particularly Z-DNA^50^. Z-DNA is a higher-energy, left-handed DNA conformation that can form under various conditions depending on sequence and environment^51^. Purine-pyrimidine repeats are particularly prone to adopting this conformation *in vitro* when stabilized by specific reagents or multivalent salts such as MgCl_2_ or hexaaminecobalt(III) chloride^52,53^, and methylation of CG repeats further stabilizes the Z-form as compared to unmethylated poly(CG)^50^. In mammalian genomes, Z-DNA-forming regions are enriched in promoters, where the structure is temporally formed due to negative supercoiling during transcription^54–56^.

**Figure 6.**
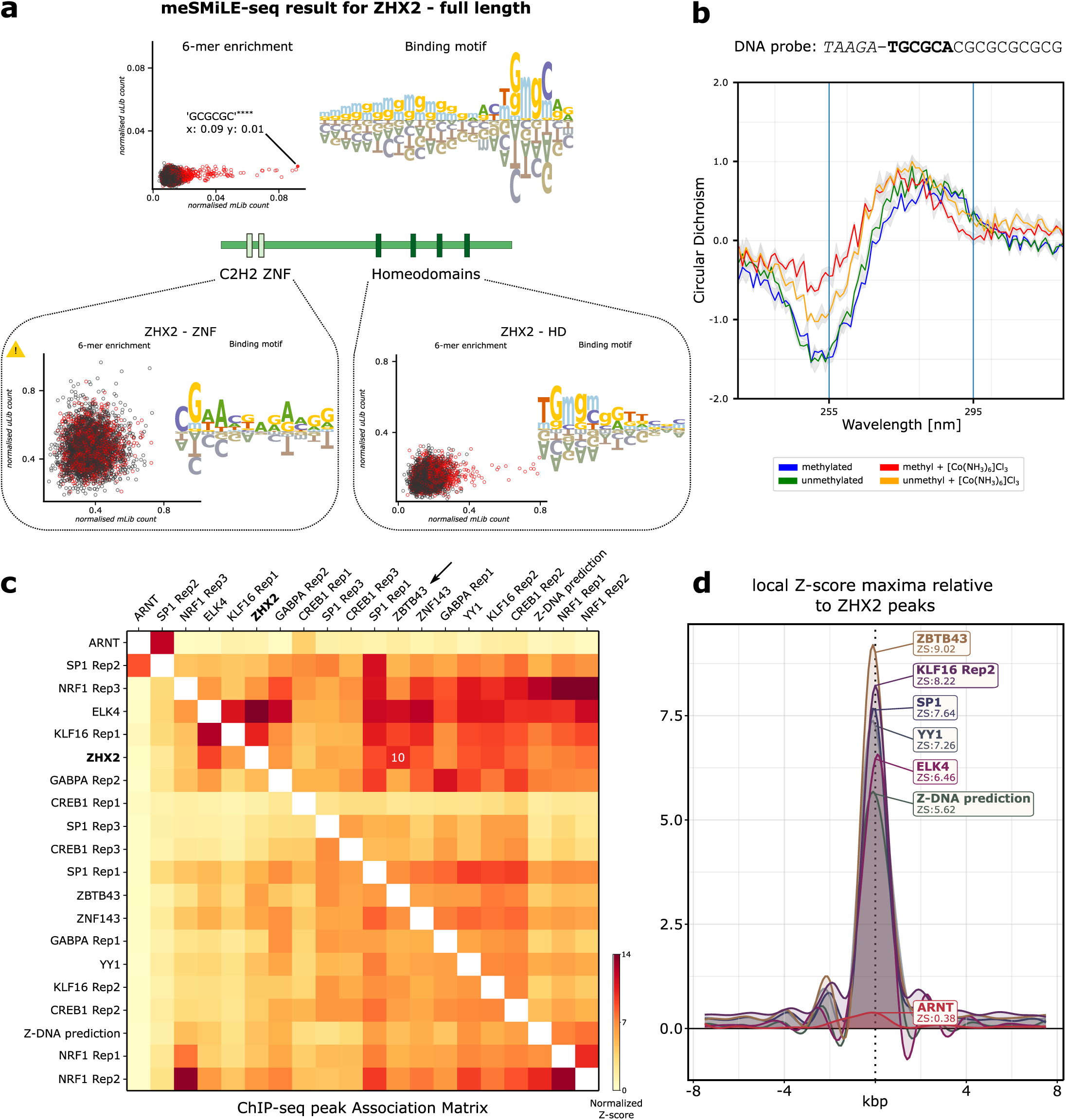
ZHX2 as a putative Z-DNA binding protein. **a)** Schematic structure of ZHX2, depicting its C2H2 ZNFs and homeodomains (HDs). The full-length protein and isolated HDs, but not the ZNFs, enrich methylated CG repeats in meSMiLE-seq, showcased by correlation scatterplots and PSAMs as described in Figure 2b**. b)** CD spectroscopy (ellipticity) of meSMiLE-seq-inferred ZHX2 binding sites plotted in function of the wavelength. The data shows B-Z transition as seen by the upshift and downshift in ellipticity at 255 nm and 295 nm, respectively. Blue and red lines: methylated DNA; green and orange lines: unmethylated DNA. Red and orange lines: added hexaaminecobalt(III) chloride. **c)** Heatmap shows ChIP-seq peak associations between TFs in HepG2 cells expressed as normalized Z-scores. ZHX2 specific peaks display highest scores with ZBTB43 peaks. **d)** Local Z-scores between selected TFs and ZHX2 within a 7.5 kbp neighborhood. The sharp drop indicates a central, peak-specific association instead of lateral, region-wide overlaps.

To test whether the meSMiLE-seq-derived sequences may form Z-DNA, thus indicating that ZHX2 might preferentially bind to this particular DNA conformation, we performed circular dichroism (CD) spectroscopy. We found that both methylated and unmethylated sequences can transition from canonical B-DNA to Z-DNA when incubated with hexaaminecobalt(III) chloride. However, methylated DNA displayed a higher potential to transition as seen by a stronger upshift in ellipticity at 255 nm (**Figure 6b**). This demonstrates that methylated ‘CG’ repeats shift the equilibrium from B-DNA towards Z-DNA compared to unmethylated sequences. The observation also suggests that Z-DNA could form in a complex medium such as the Wheat Germ IVT-kit that is used for TF production in meSMiLE-seq experiments (**Methods**).

We then searched for evidence of Z-DNA formation in the genomic target sites of ZHX2 to further test if this DNA conformation was preferentially bound by the protein. We calculated associations of ZHX2-specific regions with peaks from other TFs that were selected based on motif similarity to ZHX2 or Z-DNA from publicly available ChIP-seq data (**Methods**, **Table 5**)^55^. Additionally, we included datasets for *in silico* predicted Z-DNA forming sites^56^ and for ZBTB43, a TF that has been recently identified as a Z-DNA remodeler in prospermatogonia^57^. Most notably, ZHX2 exhibited the highest association scores and strongest local overlap with ZBTB43 across all tested TFs. These observations strongly suggest that ZHX2 is recruited to its genomic locations by recognizing Z-DNA conformations rather than canonical B-DNA motifs (**Figure 6c-d**). Altogether, our meSMiLE-seq findings suggest an alternative explanation for how ZHX2 may be recruited to promoter sequences *in vivo*. Given that Z-DNA is a transient structure in mammalian promoters, formulating this hypothesis based solely on ChIP-seq data would have been challenging. This underscores the value of *in vitro* DNA binding assays such as meSMiLE-seq.

## Discussion

In this study, we investigated the DNA binding properties of 284 putative human TFs using SMiLE-seq and 114 TFs via meSMiLE-seq, identifying motifs for 98 TFs and thus significantly expanding the TF “codebook”. One of the defining features of SMiLE-seq as a platform to profile TF-DNA interactions is the single round of DNA enrichment, as best demonstrated in meSMiLE-seq. This is because the simultaneous exposure of multiple DNA species to a TF, coupled with the single step of entrapment of bound molecules during MITOMI allows capturing the actual, potentially subtle binding preferences of TFs to modified or unmodified DNA. The lack of multiple amplification steps preserves the methylation status of the DNA and ensures the identification of TFBS over a wide range of affinities. We demonstrated that meSMiLE-seq-derived motifs correlate highly with the binding models generated by orthogonal datasets, when we excluded information about methylation specificity in the form of an extended alphabet. Since most TF-DNA interaction assays do not include modified DNA by default, there may be interest in the field to probe already characterized TFs for methylation affinity. Here, meSMiLE-seq offers a clear advantage for being a scalable competition assay that captures binding events in equilibrium conditions, as exposure to unmethylated libraries serves as an internal positive control for assessment of experimental success if a TF’s unmethylated binding site is already known. In this regard, a valuable future goal may be to expand meSMiLE-seq to include further DNA modifications which have not yet been studied to the same extent as CG methylation on potentially impacting TF-DNA interactions such as 5’-hydroxymethylation of cytosine (5hmC)^58^ or N6’-methylation of adenosine (N6mA)^59^.

The Codebook/GRECO-BIT initiative successfully identified DNA binding sites for 236 of 394 predicted human TFs. However, for 158 TFs, motif discovery did not yield reproducible motifs, suggesting a possible misclassification of proteins as TFs^1^, or a need for specific interaction partners, such as heterodimers, which were not considered within the Codebook/GRECO-BIT collaboration or in this study. Our analysis yielded 73 high-quality binding motifs from 284 TFs using SMiLE-seq, with an initial success rate of ∼26% with our analysis approach or ∼23% using non-customized motif discovery pipelines. The lower-than-expected yield was likely due to technical challenges, including a single round of DNA enrichment and bias in nucleotide distribution, which complicated motif discovery. By changing the DNA libraries and ensuring uniform nucleotide distributions in meSMiLE-seq, the success rate nearly doubled (∼42%). The improvement was achieved despite the additional experimental complexity by exposing the TFs to both methylated and unmethylated DNA, although part of this increase may be influenced by using a subset of TFs in meSMiLE-seq that was suspected to bind DNA. Nonetheless, this highlights the importance of customized analysis pipelines for overcoming background noise and extracting accurate TFBS, indicating that further optimization could significantly enhance future discoveries.

Many of our meSMiLE-seq-derived motifs could be found in TF-associated ChIP-seq peaks in HEK293/T cells. Pairing the data with WGBS revealed that the predictions for most ‘methyl minus’ TFs were correct, as motif occurrences for these TFs were predominantly not methylated in cells. Importantly, this analysis also showed that several ‘methyl plus’ TFs bound methylated genomic regions, not only indicating that binding models acquired by *in vitro* methods such as meSMiLE-seq can be translated into a cellular context, but also that these TFs are likely involved in different regulatory pathways depending on the methylation status of their motifs. For example, gene ontology enrichment analysis for ZNF18 showed that its ‘methyl plus’ binding sites were significantly associated with ‘Pathways in cancer’. This finding aligns well with DNA methylation-mediated transcriptional dysregulation, considering that aberrant DNA methylation patterns are frequent in malignancies^60,61^. The discrepancy that most ‘little effect’ and some ‘methyl plus’ TFs appeared to mostly bind unmethylated DNA in HEK293/T cells may be due to the inaccessibility of those motifs in a cell type-specific chromatin landscape since CG methylation is frequently associated with the formation of heterochromatin and gene silencing. TFs that do not possess ‘pioneering’ capability, i.e. the ability to bind condensed chromatin and/or initiate chromatin remodeling, may therefore have their TFBS occluded by nucleosomes^60,62,63^. Another constraining factor might be competition for methylated binding sites between these TFs and other TFs with high methylated DNA affinities such as MBDs, as suggested by previous studies^36^. In this case, binding to methylated motifs may be restricted to specific loci and cell types.

Lastly, our study identified ZHX2 as a potential Z-DNA binding protein, as it consistently enriched methylated purine-pyrimidine repeats. The stabilization of poly(CG) sequences in the Z-form by multivalent salts and cytosine methylation suggests that Z-DNA may be randomly adopted in meSMiLE-seq experiments where TFs are incubated with DNA for an extended time. While non-canonical DNA conformations like Z-DNA are already well known to impact gene regulation^64–66^, recent computational advancements have renewed interest in predicting genomic regions that are likely to adopt non-B-DNA structures and linking these structures to various cellular processes^56^. However, validating protein binding to these structures remains challenging due to their temporal instability under physiological conditions. In this sense, meSMiLE-seq could offer a valuable platform to more systematically study these interactions. For example, incubating proteins with DNA to reach equilibrium and adding reagents to trigger specific DNA transitions would allow the comparison of eluted DNA with and without reagents to identify interactions with specific DNA structures.

## Material and Methods

### Experimental procedures

#### TF selection and plasmids

Transcription factors were provided as EGFP fusion proteins in pF3A WG (Promega) by the Codebook/GRECO-BIT collaboration^16^. Plasmids encoding additional KRAB-Zinc finger proteins were kindly provided by Didier Trono’s laboratory and were cloned into pDONR221 plasmids (ThermoFisher Scientific) as EGFP fusions and further shuffled into custom-made Gateway-compatible pF3A WG.

#### SMiLE-seq procedure

##### 1. Library generation

meSMiLE-seq libraries comprising a random region were ordered as hand-mixed DNA oligos from IDT and resuspended to a concentration of 100 µM **(Table 4**). dsDNA libraries were synthesized via enzymatic reaction.3 µl library was mixed with 6 µl annealing_primer (100 µM), 3 µl 10x NEBuffer2 (NEB), and 18 µl ddH2O, and incubated for 5 min at 95 °C followed by 1 min at 60 °C. 20 µl were transferred into a new tube containing 16 µl ddH2O, 5 µl 10x NEBuffer2, 4 µl 10 mM dNTPs (Thermo), and 5 µl DNA Polymerase I (Large Klenow Fragment) (NEB). The reaction was incubated for 60 min at 37 °C and subsequently purified and eluted in 13 µl ddH2O using a MinElute kit (Qiagen) following the manufacturer’s instructions.

**Table 4:** DNA libraries and primers.

##### 2. Methylation of libraries for meSMiLE-seq

DNA libraries were treated differently depending on their respective mBCs (**Table 4**). uLibs remained unmodified while mLibs were methylated by mixing 250 ng of DNA with 5 µl 1.6 mM SAM (NEB), 5 µl 10x NEBuffer2 and 1 µl M.SssI (NEB) in a total volume of 50 µl (topped off with ddH2O), and incubated for 4 h at 37 °C followed by 20 min at 65 °C. To ensure a maximal degree of CG methylation, this step was performed twice. Libraries were purified using a MinElute kit (Qiagen).

The efficacy of the methylation protocol was verified by methylation of a control library (CL) (**Table 4**) followed by enzymatic digestion using BstBI (NEB). 1 μg of both modified and unmodified CLs were mixed with 1 μl enzyme, 5 μl 10x rCutSmart buffer (NEB) in a total volume of 50 μl and incubated for 15 min at 65 °C. The samples were analyzed via agarose gel electrophoresis using a 1 % agarose gel (Thermo) in 1x Tris-Acetate-EDTA buffer stained with 1:10000 SYBR Safe (Thermo).

Subsequently, mLibs and uLibs were mixed in a 1:1 molar ratio and diluted 1:10 in ddH2O. Each library aliquot of 10 µl was mixed with 50 ng of poly-dIdC (Sigma). Aliquots were stored at -20 °C for a maximum duration of 3 months.

##### 3. SMiLE-seq assay

The SMiLE-seq pipeline including chip fabrication and functionalization was carried out essentially as previously described in Isakova et al 2017^21^. In brief, TFs of interest were expressed as GFP fusion proteins using the TNT SP6 High-Yield Wheat Germ Protein Expression System from Promega (referred to as IVT-kit) following the manufacturer’s instructions. TFs were incubated with one aliquot of DNA library for at least 2 h at 25 °C and then transferred into the functionalized microfluidic device (detailed protocol Isakova *et al.*^67^), where mechanical trapping of molecular interactions was performed.

##### 4. Post-experiment library purification

Recovered DNA was purified with a MinElute kit (Qiagen) and eluted in 20 μl elution buffer. The eluate was mixed with 32.5 μl NEBNext High-Fidelity 2x PCR Master Mix (NEB), 0.5 μl library_primer_fwd (10 μM), 0.5 μl library_primer_rev (10 μM), 0.5 μl 100x SYBR Green I (Thermo) in a total reaction volume of 65 μl. 50 μl of the reaction were kept on ice while 15 μl were used to estimate the suitable amount of amplification cycles using StepOnePlus Real-Time PCR instrument (Applied Biosystems) following the subsequent program: hot start at 98 °C for 30 s and 25 cycles of 98 °C for 10 s, 63 °C for 30 s and 72 °C for 1 min, followed by a final elongation at 72 °C for 1 min, then kept at 4 °C. The remaining 50 μl were amplified accordingly and purified with a MinElute kit (Qiagen). DNA was size selected using 1.5x (one-sided) AMPureXP beads (Beckman Coutler) to remove primer dimers.

After purification, libraries were amplified for 5 cycles with 0.5 μl i5 and i7 Nextera adapters (10 μM) following the instructions provided by Illumina. Finally, DNA was purified (MinElute, Qiagen) and size selected using 1x (one-sided) AMPure XP beads (Beckman Coulter) to remove impurities.

Libraries were sequenced on Illumina MiSeq and NextSeq500 platforms. All data were acquired in four sequencing batches.

#### Electrophoretic Mobility Shift Assays

Electrophoretic mobility shift assays of PRDM13 and USF3 were performed using precast 6 % TBE gels (Novex) and were run in 0.5 % TBE buffer at 4 °C and 100 V. The TFs were expressed as GFP fusion proteins using the IVT-kit. 2 ul of unpurified IVT-TF solution was mixed with 0.1 pmol of Cy5-labeled DNA probe (IDT, **Table 4**) in 1x binding buffer (80 ng poly-dIdC, 10 mM Tris-HCl, 10 mM NaCl, 40 mM KCl, 1 mM MgCl2, 1 mM EDTA, 1 mM DTT and 0.05 mg/mL BSA). To outcompete binding between TF and labeled DNA, 1 pmol (10x molar excess) of unlabeled DNA probe (referred to as ‘cold probe’) was added where indicated. The solutions were incubated for 15 min at RT, then supplemented with 1x loading dye (0.1 M Tris-HCl, 10 % Glycerol, and 0.01 % Bromophenol blue (Sigma)) and loaded onto the gel. After electrophoresis, gels were imaged using an Amersham Typhoon scanner (Cytiva).

#### CD spectroscopy

CD spectroscopy measurements were conducted on a Chirascan V100 (AppliedPhotophysics) using 10 μM DNA probes (Merck) in 270 μl (**Table 4**) CD buffer (15 mM NaCl, 10 mM Tris-HCl) with and without hexaaminecobalt(III) chloride (1 mM) (Sigma). DNA probes were incubated at 37 °C for 2 hours to allow a potential B-Z transition. CD measurements were acquired between 230 and 320 nm with a bandwidth of 1 nm and intervals averaged over 0.5 s at 25 °C.

### Data analysis

#### SMiLE-seq analysis

Sequenced reads were filtered and demultiplexed using custom Python scripts available on GitHub (https://github.com/DeplanckeLab/meSMiLEseq) (version 3.9.5) and pandas (version 2.0.3)^68,69^. To test for enrichment of DNA motifs, the random stretches from both input and eluted libraries were split into *k*-mers (by default 6, 7, 8, and 9-mers). *K*-mers from both input and eluted fractions were counted and used to assess significantly enriched *k*-mers in the eluted libraries using a right-tailed, one-sided Fisher’s exact test. Calculated p-values were corrected for multiple testing via the Benjamini-Hochberg method (p-value threshold 0.05). Raw sequencing reads without significantly enriched *k*-mers were filtered out; de novo motif discovery was performed with the ProBound Suite^23^ as described below. All calculations were performed using NumPy (version 1.26.4) and SciPy (version 1.13.1)^70,71^.

#### meSMiLE-seq analysis

Data were processed and analyzed as described above (SMiLE-seq analysis) for both uLib and mLib data to assess enrichment of unmethylated and methylated *k*-mers. Scatterplots were created using the ratios of respective *k*-mers (eluted count/ input count) considering their methylation status unless otherwise stated. Empirical cumulative distribution functions (ECDFs) were plotted using Z-transformed *k*-mer counts from the input libraries. Graphs were plotted using matplotlib (version 3.6.2)^72^.

Motif similarities were assessed by flattening both PSAMs and PFMs into one-dimensional arrays. Pearson correlation coefficients (PCC) were then calculated between these flattened vectors for each TF pair. For motifs of unequal lengths, the shorter motif was used as a sliding window across the longer motif, and all possible PCCs were computed. The most extreme PCC value (either closest to 1 (indicating high correlation) or -1 (indicating high anticorrelation)) was reported.

DNA motifs were visualized using the Python package logomaker^73^.

#### ProBound analysis

SMiLE-seq and meSMiLE-seq data were passed to the ProBound Suite^23^ for de novo motif discovery using the same optimizer settings that were used for ProBound benchmarking in MEX (**MEX paper**): i.e., the lambdaL2 parameter was set to 0.000001, Dirichlet regularizer weight to 20 and the likelihoodThreshold parameter to 0.000218. meSMiLE-seq data were analyzed using ProBound’s methylation-aware binding models with an extended alphabet, where ‘mg’ and ‘CG’ indicated methylated and ‘naked’ CG dinucleotides, respectively. Data were given to ProBound as full sequences, with the input libraries serving as background. Binding affinities were modeled for both specific and non-specific binding (in total three binding modes), using sizes of 6, 9, 12, 15, and 24 base pairs.

To compare PSAM models to PFMs from other datasets, PSAMs were converted into PFMs as described in **Supplementary Figure 4a**. To extract ‘methyl plus’ motifs, PSAMs were manually adapted by swapping values of ‘mg’ dinucleotides with ‘CGs’ at respective positions. Similarly, for ‘methyl minus’ or ‘no CG’ motifs values for ‘m’ and ‘g’ nucleotides were not considered when converting PSAMs into PFMs.

#### HOMER analysis

MeSMiLE-seq sequences were split according to the methylation status using the mBC. Datasets of eluted and input libraries were stored as .fasta-files and analyzed by calling the program ‘findmotifs.pl’ with the parameters set to human, -fastaBg and -len 6, 8, 10, 12, using the input libraries as background^24^.

Data for KRAB-ZNF that were not part of the Codebook/GRECO-BIT consortium data (Geo accession number GSE78099)^29^ (**Table 5**) were analyzed by calling ‘findmotifsGenome.pl’ using hg19 as reference genome and a search window of 200 bp.

**Table 5:** Orthogonal datasets used in this study.

#### TF classification

TFs were classified based on ProBound generated PSAM models, where letter sizes represent the impact of a nucleotide on the stability of the TF-DNA interaction. If a reported motif contained a ‘CG’ and no ‘mg’ dinucleotide at a given position, and the values of both ‘C’ and ‘G‘ were > 10 % of the Euclidean norm of the PSAM at this position, the TF would be classified as ‘methyl minus’. Vice versa, if ‘mg’ was reported instead of ‘CG’, the TF would be classified as ‘methyl plus’. If both ‘CG’ and ‘mg’ were present and the condition mentioned above was met, the ratio ‘CG’/’mg’ was calculated. If values fluctuated by 10 % or more, the TFs would be classified in the respective groups.

In case none of the above was met, the TF would be labeled as ‘little effect/no CG dinucleotide’.

#### Motif occurrences in ChIP-seq and ChIP-exo data and their methylation status

All ChIP-seq and ChIP-exo datasets were mapped to or lifted over to the reference genome hg19. Sequences were extracted from TF-specific datasets (**Table 5**) by calling ‘bedtools getfasta’^74^. Methylation-sensitive PSAM models were transformed into PFMs in MEME format as described above, and all matrices were elongated by the background model used in the FIMO tool from the MEME suite to have a standard size of 20 bp^26,75^. To find individual motif occurrences within TF-specific peaks, PFMs were applied using FIMO with default parameters (p-value threshold < 10-4). The resulting file ‘best_site.narrowPeak’ was intersected with publicly available WGBS data for HEK293/T cells^34,35^ (**Table 5**) via ‘bedtools intersect’^74^ to obtain information about DNA methylation. Motif-specific methylation patterns were compared to overall CG methylation levels of peaks containing those motifs. WGBS data were filtered for lowly covered regions to ensure the recommended average coverage of 15x^76^. Significance of motif-specific methylation distributions was assessed by performing a Kolmogorov-Smirnov test with the background methylation distribution.

In addition, genomic loci were intersected with ChromHMM tracks for HEK293/T cells^37^ to extract motif-specific chromatin annotations. Datasets for ‘methyl plus’ TFs were split based on methylation levels, comparing motif occurrences being at least 50% methylated to those below the threshold with adjusted number of reads. These TFBS containing regions were used for methylation-specific gene ontology (GO) enrichment using Enrichr within a 3 kb distance of the motif^77–79^.

#### Calculating ChIP-seq associations for ZHX2

Permutation-based global associations between ChIP-seq tracks (**Table 5**) were calculated with the regioneReloaded package in R (version 4.3.1)^80^ using the ‘crosswisePermTest’ function with the settings ‘resampleGenome’, ntimes = 1000, evFUN = ‘numOverlaps’. Local associations were calculated with the ‘multiLocalZscore’ function with the same settings mentioned above, a sliding window of 7500 bp and a step size of 100 bp.

**Database 1: Overview of all meSMiLE-seq-derived DNA binding motifs**

**Database 2: Methylation levels of genomic TFBS**

## Supporting information

Supplementary Database 1

Supplementary Database 2

## Data and Code availability

Raw SMiLE-seq sequencing datasets were deposited on ArrayExpress (meSMS data: E-MTAB-14597; SMS data: E-MTAB-14598). Custom analysis pipelines and PSAMs with logos used in this manuscript are available on Github (https://github.com/DeplanckeLab/meSMiLEseq). Additionally, motifs can be browsed at mex.autosome.org, https://cisbp.ccbr.utoronto.ca/. An updated list of human TFs is available at https://humantfs.ccbr.utoronto.ca/. Information on constructs, experiments, analyses, processed data, comparison tracks, and browsable pages with information and results for each TF is available at codebook.ccbr.utoronto.ca.

## Author contributions

A.G. and B.D. designed the study. A.G., A.Y. and C.O. conducted experiments. A.G., K.F. and J.K. performed data analyses. C.S., J.R. and N.G. manufactured microfluidic devices. A.G., G.v.M. and B.D. wrote the manuscript with support from I.K., T.H., J.K and other members of the Codebook/GRECO-BIT Consortium.

## Acknowledgements

We extend our gratitude to the members of the Deplancke laboratory for their valuable input on the experiments and analyses, with special thanks to V. Gardeux and C. Lambert. Additionally, we thank the gene expression core facility and the protein production and structure core facility at the École Polytechnique Fédérale de Lausanne (EPFL) for their assistance in library sequencing and CD spectroscopy.

## The Codebook / GRECO-BIT Consortium

**Principal investigators (steering committee)**

Philipp Bucher, Bart Deplancke, Oriol Fornes, Jan Grau, Ivo Grosse, Timothy R. Hughes, Arttu Jolma, Fedor A. Kolpakov, Ivan V. Kulakovskiy, Vsevolod J. Makeev

**Analysis Centers:**

**University of Toronto (Data production and analysis):** Mihai Albu, Marjan Barazandeh, Alexander Brechalov, Zhenfeng Deng, Ali Fathi, Arttu Jolma, Chun Hu, Timothy R. Hughes, Samuel A. Lambert, Kaitlin U. Laverty, Zain M. Patel, Sara E. Pour, Rozita Razavi, Mikhail Salnikov, Ally W.H. Yang, Isaac Yellan, Hong Zheng

**Institute of Protein Research (Data analysis):** Ivan V. Kulakovskiy, Georgy Meshcheryakov

**EPFL, École polytechnique fédérale de Lausanne (Data production and analysis):** Giovanna Ambrosini, Bart Deplancke, Antoni J. Gralak, Sachi Inukai, Judith F. Kribelbauer-Swietek

**Martin Luther University Halle-Wittenberg (Data analysis):** Jan Grau, Ivo Grosse, Marie-Luise Plescher

**Sirius University of Science and Technology (Data analysis):** Semyon Kolmykov, Fedor Kolpakov

**Biosoft.Ru (Data analysis):** Ivan Yevshin

**Faculty of Bioengineering and Bioinformatics, Lomonosov Moscow State University (Data analysis):** Nikita Gryzunov, Ivan Kozin, Mikhail Nikonov, Vladimir Nozdrin, Arsenii Zinkevich

**Institute of Organic Chemistry and Biochemistry (Data analysis):** Katerina Faltejskova

**Max Planck Institute of Biochemistry (Data analysis):** Pavel Kravchenko

**Swiss Institute for Bioinformatics (Data analysis):** Philipp Bucher

**University of British Columbia (Data analysis):** Oriol Fornes

**Vavilov Institute of General Genetics (Data analysis):** Sergey Abramov, Alexandr Boytsov, Vasilii Kamenets, Vsevolod J. Makeev, Dmitry Penzar, Anton Vlasov, Ilya E. Vorontsov

**McGill University (Data analysis):** Aldo Hernandez-Corchado, Hamed S. Najafabadi

**Memorial Sloan Kettering (Data production and analysis):** Kaitlin U. Laverty, Quaid Morris

**Cincinnati Children’s Hospital (Data analysis):** Xiaoting Chen, Matthew T. Weirauch

## The Codebook / GRECO-BIT Consortium - Acknowledgments

We thank the IT Group of the Institute of Computer Science at Halle University for computational resources, Maximilian Biermann for valuable technical support, Gherman Novakovsky for providing feedback, Berat Dogan for testing earlier versions of RCADEEM, and Debashish Ray for assistance with database depositions.

This work was supported by the following:

- Canadian Institutes of Health Research (CIHR) grants FDN-148403, PJT-186136, PJT-191768, and PJT-191802, and NIH grant R21HG012258 to T.R.H.
- CIHR grant PJT-191802 to T.R.H. and H.S.N.
- Natural Sciences and Engineering Research Council of Canada (NSERC) grant RGPIN-2018-05962 to H.S.N.
- Russian Science Foundation grant 20-74-10075 to I.V.K.
- Russian Science Foundation grant 24-14-20031 to F.A.K.
- Swiss National Science Foundation grant (no. 310030_197082) to B.D.
- Marie Skłodowska-Curie (no. 895426) and EMBO long-term (1139-2019) fellowships to J.F.K.
- NIH grants R01HG013328 and U24HG013078 to M.T.W., T.R.H., and Q.M.
- NIH grants R01AR073228, P30AR070549, and R01AI173314 to M.T.W.
- NIH grant P30CA008748 partially supported Q.M.
- Canada Research Chairs funded by CIHR to T.R.H. and H.S.N.
- Ontario Graduate Scholarships to K.U.L and I.Y.
- A.J. was supported by Vetenskapsrådet (Swedish Research Council) Postdoctoral Fellowship (2016-00158)
- The Billes Chair of Medical Research at the University of Toronto to T.R.H.
- EPFL Center for Imaging
- Institutional funding from EPFL
- Resource allocations from the Digital Research Alliance of Canada

