## Supplementary Figures for "Identification of methylation-sensitive human transcription factors using meSMiLE-seq"

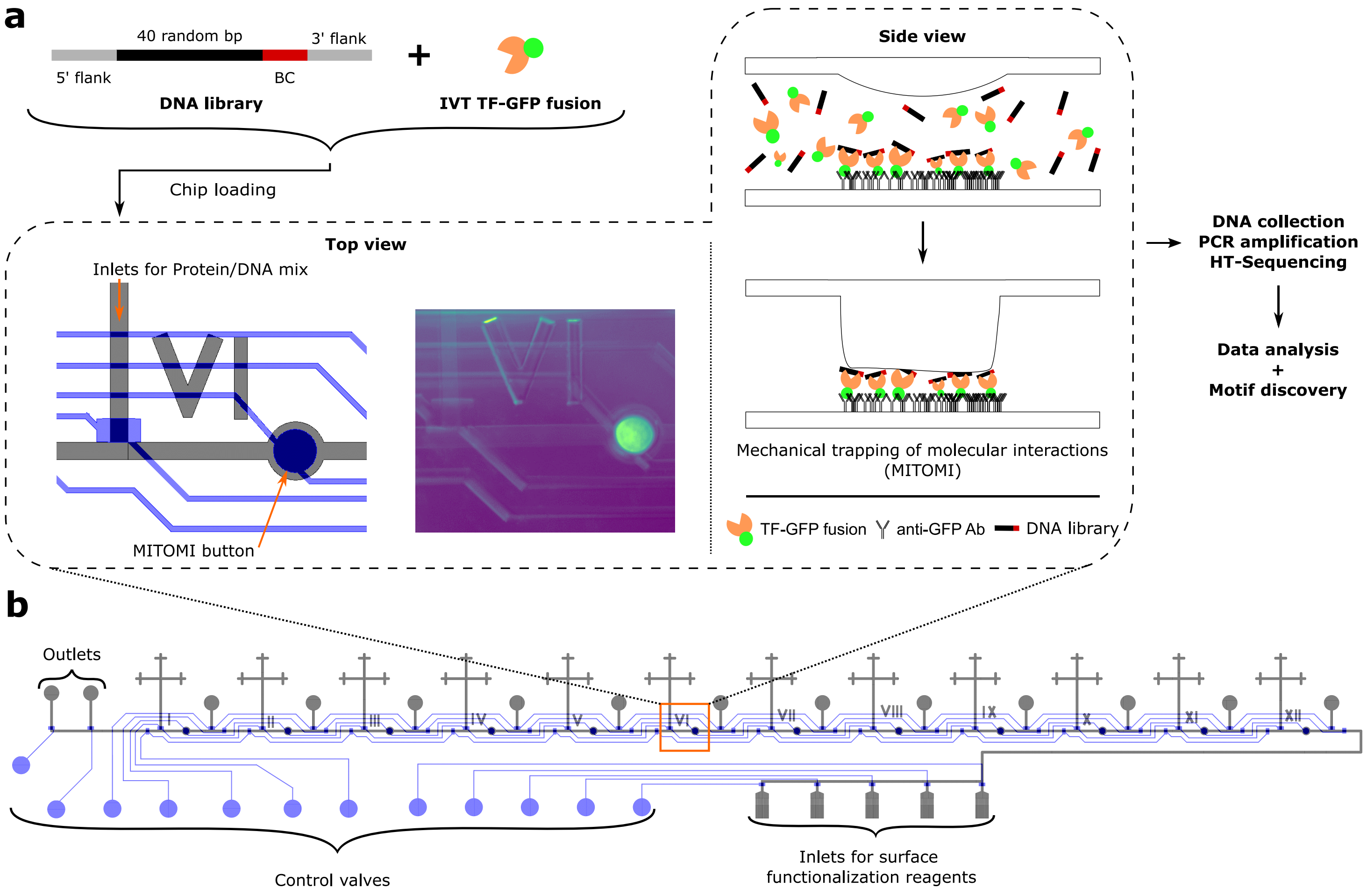

a
40 random bp
3' flank
5' flank
BC
DNA library
IVT TF-GFP fusion
Chip loading
Side view
Top view
Inlets for Protein/DNA mix
MITOMI button
Mechanical trapping of molecular interactions (MITOMI)
TF-GFP fusion
anti-GFP Ab
DNA library
DNA collection
PCR amplification
HT-Sequencing
Data analysis
+
Motif discovery
b
Outlets
Control valves
Inlets for surface functionalization reagents

**Supplementary Figure 1.**

**a)** Illustration of the SMiLE-seq workflow with schematic description of DNA libraries used in classical SMiLE-seq experiments. BC: Barcode, IVT: *in vitro* transcription and translation. **b)** Schematic structure of a SMiLE-seq chip, the control layer is depicted in blue, the flow layer in black. Each chip contains 12 MITOMI buttons allowing for 12 separate experiments per assay.

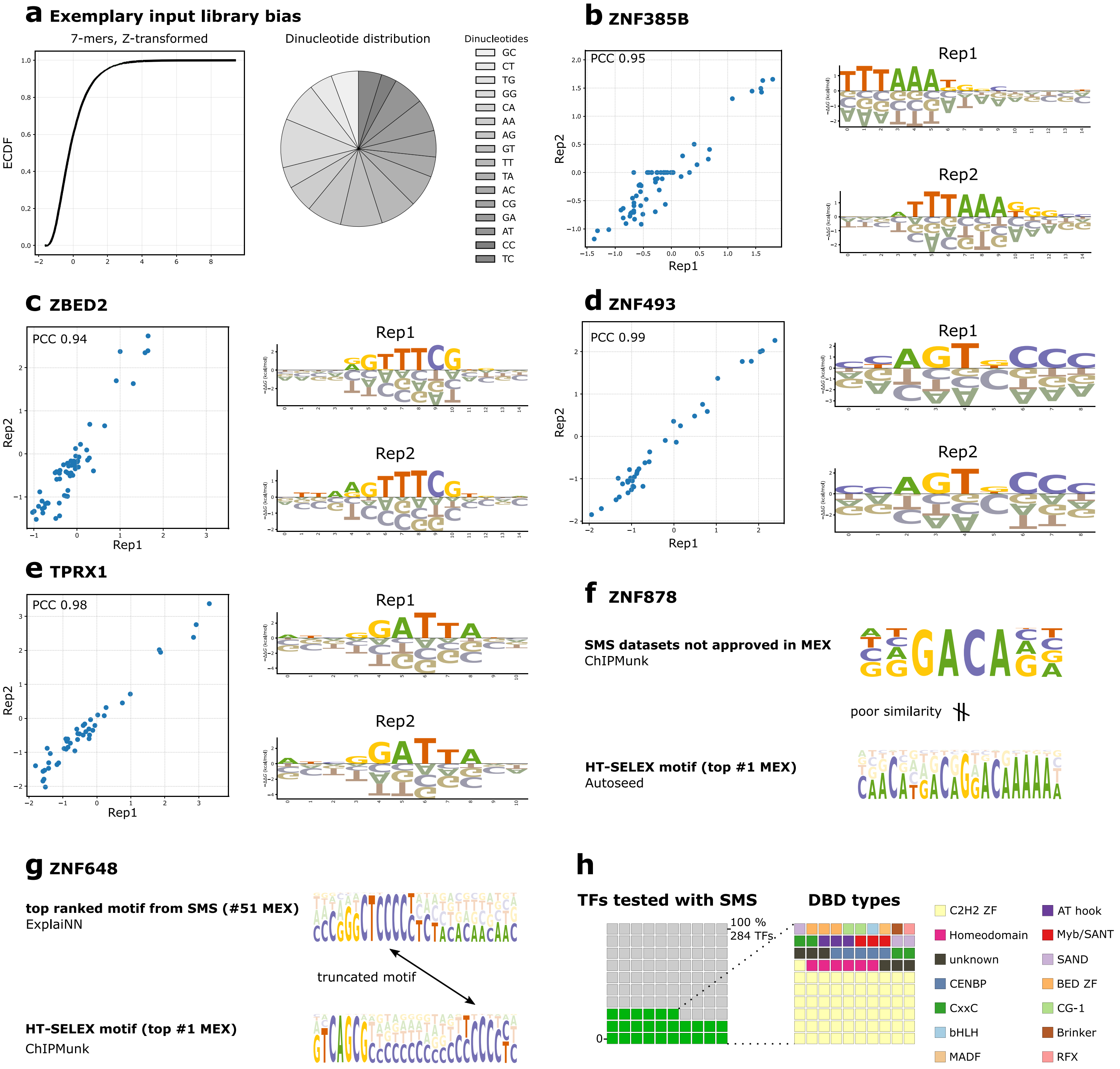
a Exemplary input library bias
7-mers, Z-transformed
ECDF
Dinucleotide distribution
Dinucleotides
GC
CT
TG
GG
CA
AA
AG
GT
TT
TA
AC
CG
GA
AT
CC
TC
b ZNF385B
PCC 0.95
Rep1
Rep2
c ZBED2
PCC 0.94
d ZNF493
PCC 0.99
e TPRX1
PCC 0.98
f ZNF878
SMS datasets not approved in MEX
ChIPMunk
poor similarity
HT-SELEX motif (top #1 MEX)
Autoseed
g ZNF648
top ranked motif from SMS (#51 MEX)
ExplaiNN
truncated motif
HT-SELEX motif (top #1 MEX)
ChIPMunk
h
TFs tested with SMS
DBD types
100 %
284 TFs
0
C2H2 ZF
Homeodomain
unknown
CENBP
CxxC
bHLH
MADF
AT hook
Myb/SANT
SAND
BED ZF
CG-1
Brinker
RFX

**Supplementary Figure 2.**

**a)** Empirical cumulative distribution function (ECDF) of 7-mers and the dinucleotide distribution of an exemplary input library used in classical SMiLE-seq. **b) to e)** Correlation scatterplots of flattened, ProBound generated PSAMs for ZNF385B, ZBED2, ZNF493, and TPRX1. PCC: Pearson correlation coefficient. **f) and g)** PFMs inferred from SMiLE-seq data using non-customized standard motif discovery tools compared to top ranked motifs in MEX for ZNF878 and ZNF648. **h)** Distribution of TF families that yielded DNA motifs in this study.

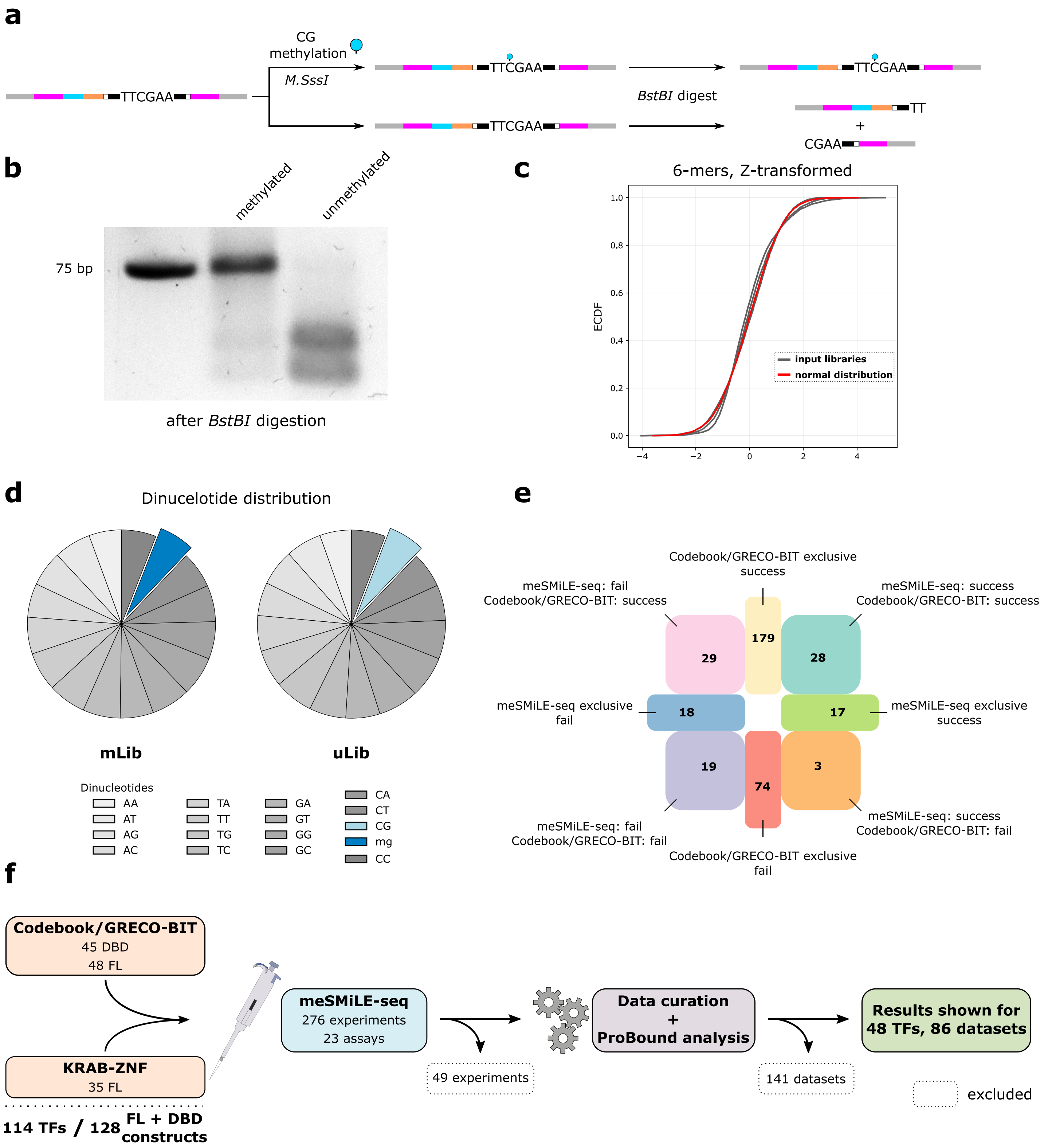
a
CG
methylation
M.SssI
TTCGAA
BstBI digest
TT
+
CGAA
b
methylated
unmethylated
75 bp
after BstBI digestion
c
6-mers, Z-transformed
ECDF
input libraries
normal distribution
d
Dinucelotide distribution
mLib
uLib
Dinucleotides
AA
AT
AG
AC
TA
TT
TG
TC
GA
GT
GG
GC
CA
CT
CG
mg
CC
e
Codebook/GRECO-BIT exclusive success
meSMiLE-seq: fail Codebook/GRECO-BIT: success
meSMiLE-seq: success Codebook/GRECO-BIT: success
29
179
28
meSMiLE-seq exclusive fail
18
17
meSMiLE-seq exclusive success
19
74
3
meSMiLE-seq: fail Codebook/GRECO-BIT: fail
meSMiLE-seq: success Codebook/GRECO-BIT: fail
Codebook/GRECO-BIT exclusive fail
f
Codebook/GRECO-BIT
45 DBD
48 FL
KRAB-ZNF
35 FL
114 TFs / 128 FL + DBD constructs
meSMiLE-seq
276 experiments
23 assays
49 experiments
Data curation
+
ProBound analysis
141 datasets
Results shown for 48 TFs, 86 datasets
excluded

**Supplementary Figure 3.**

**a)** Control library designed to optimize the methylation reaction. The structure is equivalent to other meSMiLE-seq libraries, with an additional cleavage site for the methylation-sensitive restriction enzyme *BstBI* within the random region. **b)** Agarose gel electrophoresis of methylated and unmethylated control libraries after *BstBI* digest. **c) and d)** ECDF of 6-mers and the dinucleotide distribution of newly designed DNA libraries used in meSMiLE-seq. **e)** Overlap of TFs assayed in meSMiLE-seq and the Codebook/GRECO-BIT collaboration. Numbers indicate the number of TFs that belong to the reported description. For example, 'meSMiLE-seq exclusive success/fail' indicates the number of TFs that were assayed solely via meSMiLE-seq and not in the collaboration, out of which 17 yielded DNA motifs and 18 did not. **f)** Schematic overview of how many TFs were assayed with meSMiLE-seq and for how many TFs, DNA binding motifs were approved.

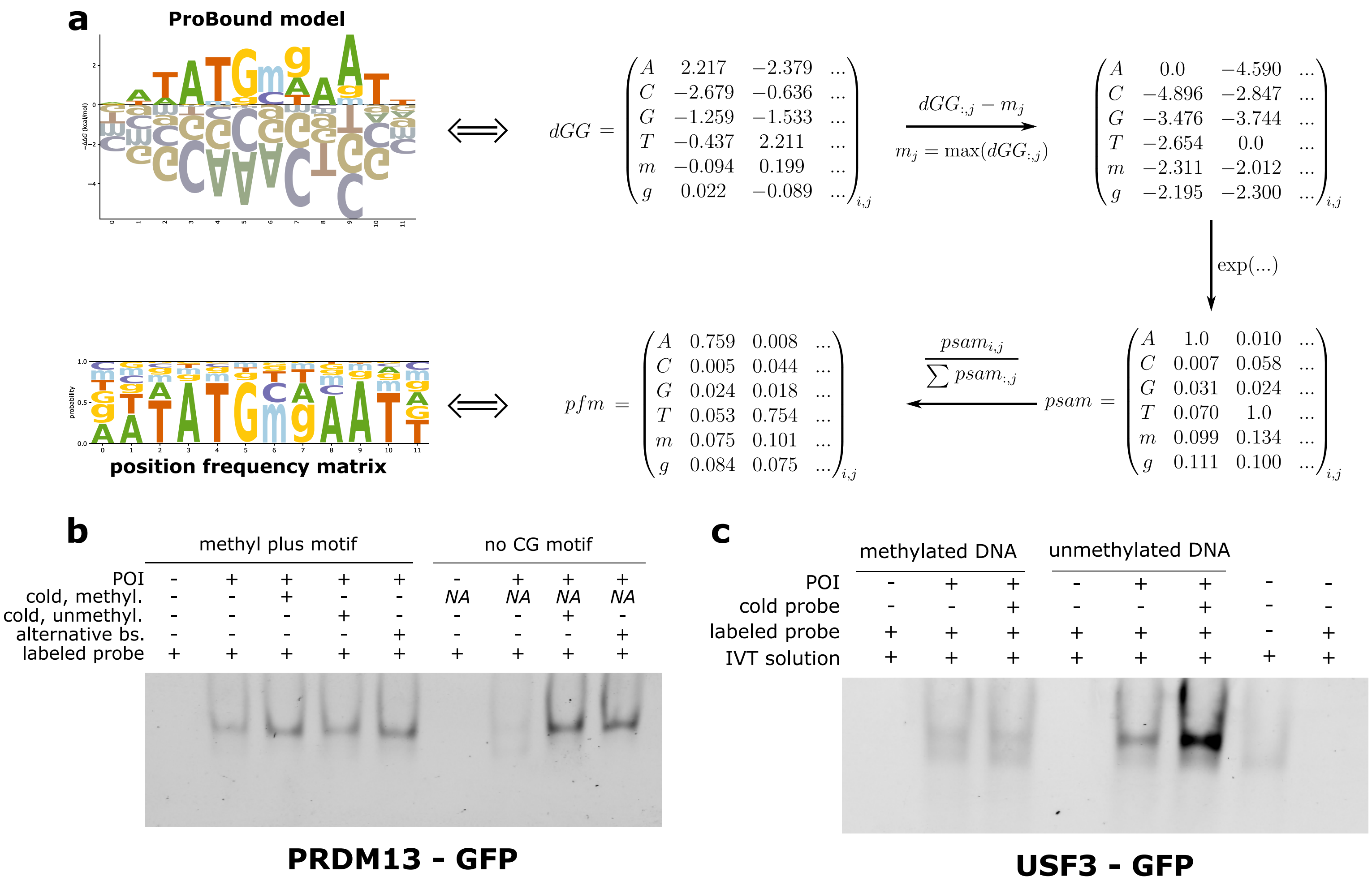
a
ProBound model
position frequency matrix
dGG =
A 2.217 −2.379 ...
C −2.679 −0.636 ...
G −1.259 −1.533 ...
T −0.437 2.211 ...
m −0.094 0.199 ...
g 0.022 −0.089 ...
dGG:,j − mj
mj = max(dGG:,j)
A 0.0 −4.590 ...
C −4.896 −2.847 ...
G −3.476 −3.744 ...
T −2.654 0.0 ...
m −2.311 −2.012 ...
g −2.195 −2.300 ...
exp(...)
psam =
A 1.0 0.010 ...
C 0.007 0.058 ...
G 0.031 0.024 ...
T 0.070 1.0 ...
m 0.099 0.134 ...
g 0.111 0.100 ...
psami,j / ∑ psam:,j
pfm =
A 0.759 0.008 ...
C 0.005 0.044 ...
G 0.024 0.018 ...
T 0.053 0.754 ...
m 0.075 0.101 ...
g 0.084 0.075 ...
b
methyl plus motif
no CG motif
POI
cold, methyl.
cold, unmethyl.
alternative bs.
labeled probe
NA
PRDM13 - GFP
c
methylated DNA
unmethylated DNA
POI
cold probe
labeled probe
IVT solution
USF3 - GFP

**Supplementary Figure 4.**

**a)** Schematic description on how to transform PSAM models into PFMs. To exclude information about methylation sensitivity, ‘m’ and ‘g’ rows are omitted. **b) and c)** EMSA gels of PRDM13 and USF3 as reported in **Figure 4b-c**. Here, the signal corresponds to TF-GFP fusion proteins instead of labeled DNA probes as shown in the main figure. For the legend, see **Figure 4**.

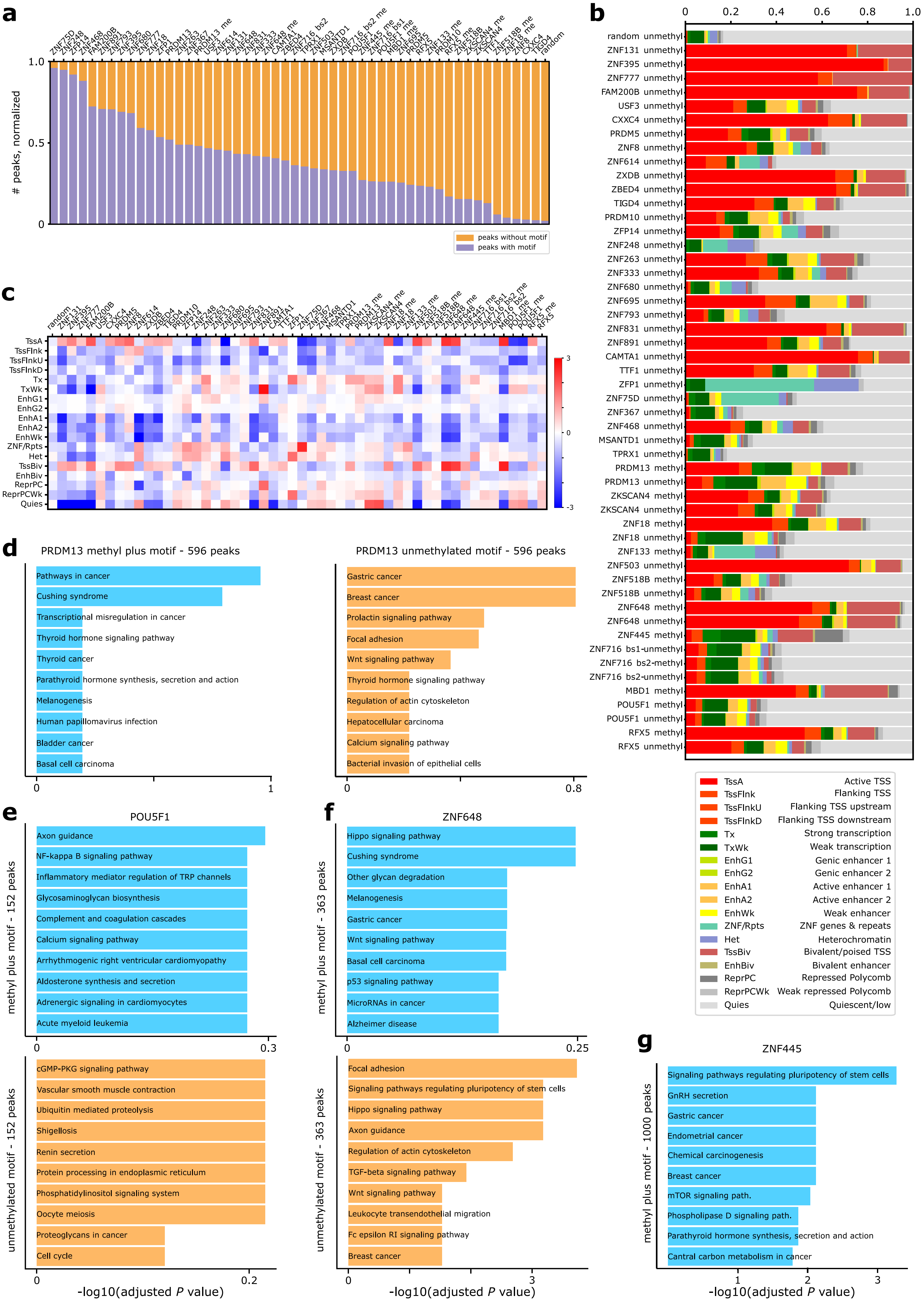

a
### peaks, normalized
1.0
0.5
0
peaks without motif
peaks with motif
b
0
0.2
0.4
0.6
0.8
1.0
random unmethyl
ZNF131 unmethyl
ZNF395 unmethyl
ZNF777 unmethyl
FAM200B unmethyl
USF3 unmethyl
CXXC4 unmethyl
PRDM5 unmethyl
ZNF8 unmethyl
ZNF614 unmethyl
ZXDB unmethyl
ZBED4 unmethyl
TIGD4 unmethyl
PRDM10 unmethyl
ZFP14 unmethyl
ZNF248 unmethyl
ZNF263 unmethyl
ZNF333 unmethyl
ZNF680 unmethyl
ZNF695 unmethyl
ZNF793 unmethyl
ZNF831 unmethyl
ZNF891 unmethyl
CAMTA1 unmethyl
TTF1 unmethyl
ZFP1 unmethyl
ZNF75D unmethyl
ZNF367 unmethyl
ZNF468 unmethyl
MSANTD1 unmethyl
TPRX1 unmethyl
PRDM13 methyl
PRDM13 unmethyl
ZKSCAN4 methyl
ZKSCAN4 unmethyl
ZNF18 methyl
ZNF18 unmethyl
ZNF133 methyl
ZNF503 unmethyl
ZNF518B methyl
ZNF518B unmethyl
ZNF648 methyl
ZNF648 unmethyl
ZNF445 methyl
ZNF716 bs1-unmethyl
ZNF716 bs2-methyl
ZNF716 bs2-unmethyl
MBD1 methyl
POU5F1 methyl
POU5F1 unmethyl
RFX5 methyl
RFX5 unmethyl
TssA Active TSS
TssFlnk Flanking TSS
TssFlnkU Flanking TSS upstream
TssFlnkD Flanking TSS downstream
Tx Strong transcription
TxWk Weak transcription
EnhG1 Genic enhancer 1
EnhG2 Genic enhancer 2
EnhA1 Active enhancer 1
EnhA2 Active enhancer 2
EnhWk Weak enhancer
ZNF/Rpts ZNF genes & repeats
Het Heterochromatin
TssBiv Bivalent/poised TSS
EnhBiv Bivalent enhancer
ReprPC Repressed Polycomb
ReprPCWk Weak repressed Polycomb
Quies Quiescent/low
c
TssA
TssFlnk
TssFlnkU
TssFlnkD
Tx
TxWk
EnhG1
EnhG2
EnhA1
EnhA2
EnhWk
ZNF/Rpts
Het
TssBiv
EnhBiv
ReprPC
ReprPCWk
Quies
3
0
-3
d
PRDM13 methyl plus motif - 596 peaks
Pathways in cancer
Cushing syndrome
Transcriptional misregulation in cancer
Thyroid hormone signaling pathway
Thyroid cancer
Parathyroid hormone synthesis, secretion and action
Melanogenesis
Human papillomavirus infection
Bladder cancer
Basal cell carcinoma
0
1
PRDM13 unmethylated motif - 596 peaks
Gastric cancer
Breast cancer
Prolactin signaling pathway
Focal adhesion
Wnt signaling pathway
Thyroid hormone signaling pathway
Regulation of actin cytoskeleton
Hepatocellular carcinoma
Calcium signaling pathway
Bacterial invasion of epithelial cells
0
0.8
e
POU5F1
methyl plus motif - 152 peaks
Axon guidance
NF-kappa B signaling pathway
Inflammatory mediator regulation of TRP channels
Glycosaminoglycan biosynthesis
Complement and coagulation cascades
Calcium signaling pathway
Arrhythmogenic right ventricular cardiomyopathy
Aldosterone synthesis and secretion
Adrenergic signaling in cardiomyocytes
Acute myeloid leukemia
0
0.3
unmethylated motif - 152 peaks
cGMP-PKG signaling pathway
Vascular smooth muscle contraction
Ubiquitin mediated proteolysis
Shigellosis
Renin secretion
Protein processing in endoplasmic reticulum
Phosphatidylinositol signaling system
Oocyte meiosis
Proteoglycans in cancer
Cell cycle
0
0.2
-log10(adjusted P value)
f
ZNF648
methyl plus motif - 363 peaks
Hippo signaling pathway
Cushing syndrome
Other glycan degradation
Melanogenesis
Gastric cancer
Wnt signaling pathway
Basal cell carcinoma
p53 signaling pathway
MicroRNAs in cancer
Alzheimer disease
0
0.25
unmethylated motif - 363 peaks
Focal adhesion
Signaling pathways regulating pluripotency of stem cells
Hippo signaling pathway
Axon guidance
Regulation of actin cytoskeleton
TGF-beta signaling pathway
Wnt signaling pathway
Leukocyte transendothelial migration
Fc epsilon RI signaling pathway
Breast cancer
0
3
-log10(adjusted P value)
g
ZNF445
methyl plus motif - 1000 peaks
Signaling pathways regulating pluripotency of stem cells
GnRH secretion
Gastric cancer
Endometrial cancer
Chemical carcinogenesis
Breast cancer
mTOR signaling path.
Phospholipase D signaling path.
Parathyroid hormone synthesis, secretion and action
Cantral carbon metabolism in cancer
1
2
3
-log10(adjusted P value)

**Supplementary Figure 5.**

**a)** Bar plot depicts percentages of TF-specific ChIP-seq peaks in which meSMiLE-seq motifs were found, using the method shown in **Figure 5a-b. b) and c)** ChromHMM annotations of individual motif occurrences in TF-specific ChIP-seq peaks for all TFs, depicted as bar charts and as log2-transformed ratios as described in **Figure 5e-f. d) to g)** Gene ontology enrichment analysis of 'methyl plus' TFs PRDM13, POU5F1, ZBF648 and ZNF445.
