## Supplementary Database 1 for "Identification of methylation-sensitive human transcription factors using meSMiLE-seq"

⚠ = weak enrichment

### C2H2 ZNF

#### PRDM5 DBD: weak methyl minus

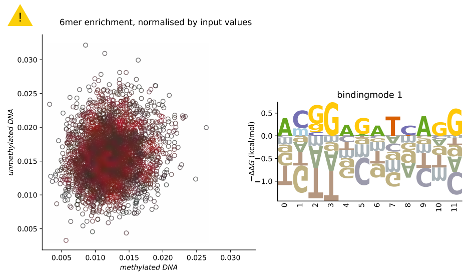


#### PRDM10 DBD: little effect

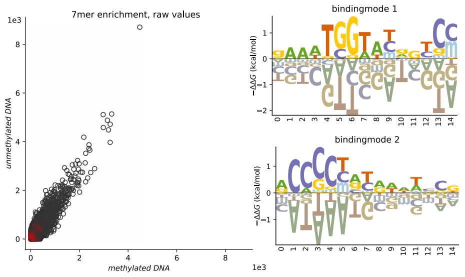


#### PRDM13 DBD: weak methyl plus

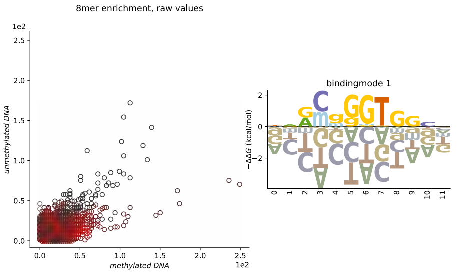


#### ZBTB5 DBD: no CG

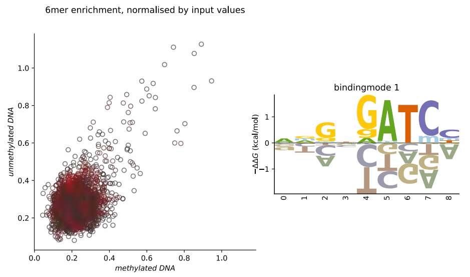


#### ZBTB46 DBD: little effect

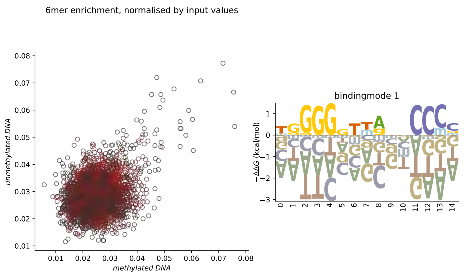


#### ZFP1 FL: no CG

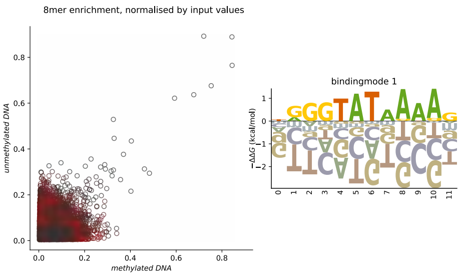


#### ZFP14 FL: little effect

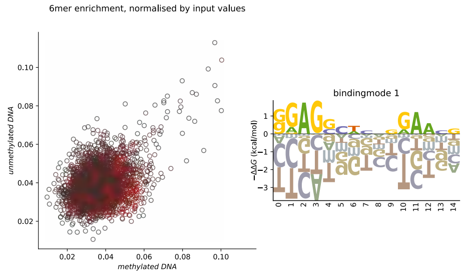


#### ZKSCAN4 DBD: weak methyl plus

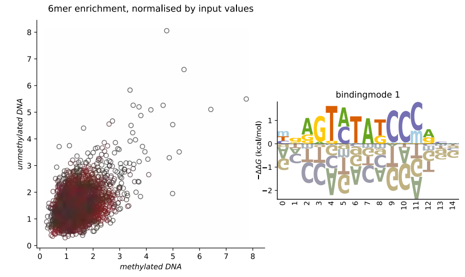


#### ZNF8 FL: weak methyl minus

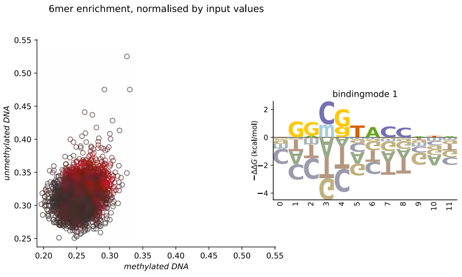


#### ZNF18 FL: weak methyl plus

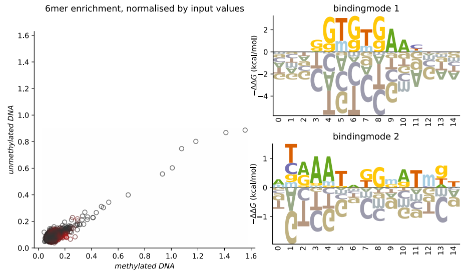


#### ZNF23 FL: weak methyl plus

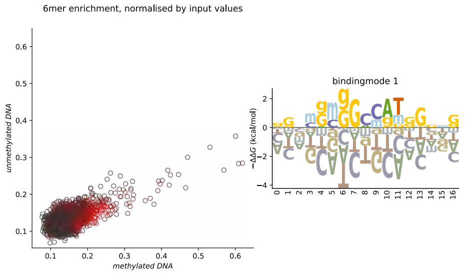


#### ZNF75D FL: no CG

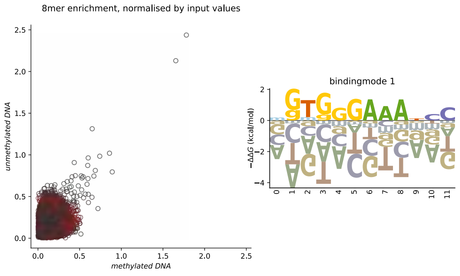


#### ZNF131 FL: methyl minus

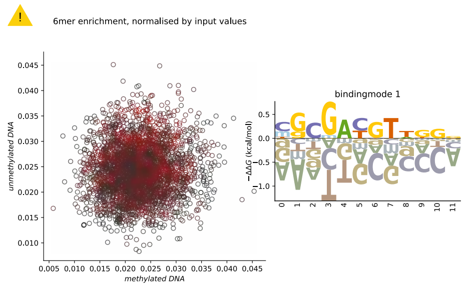


#### ZNF133 FL: weak methyl plus

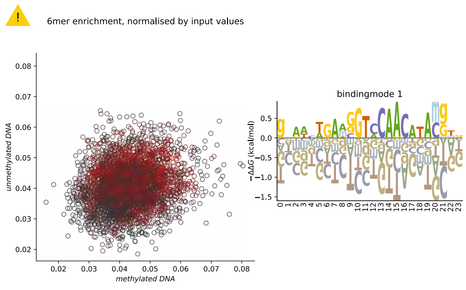


#### ZNF248 FL: little effect

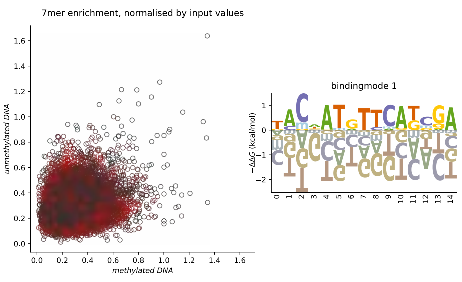

### C2H2 ZNF

##### ZNF263 FL: little effect

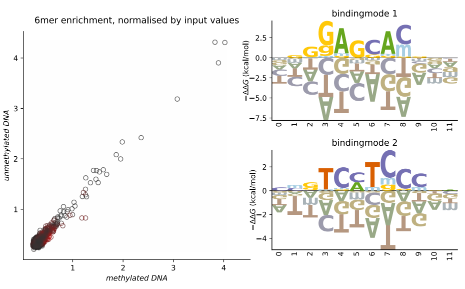


##### ZNF333 FL: little effect

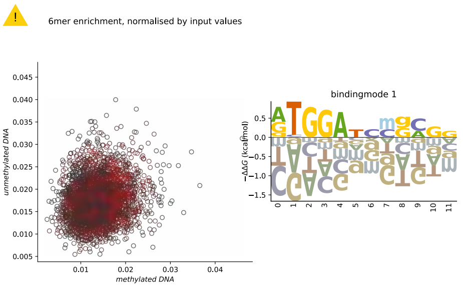


##### ZNF367 FL: no CG

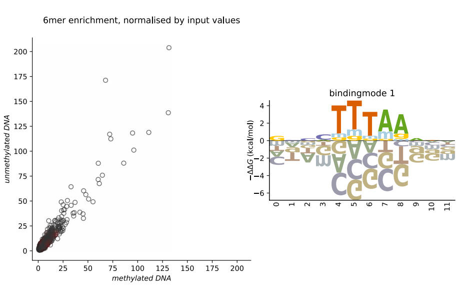


##### ZNF395 DBD: methyl minus

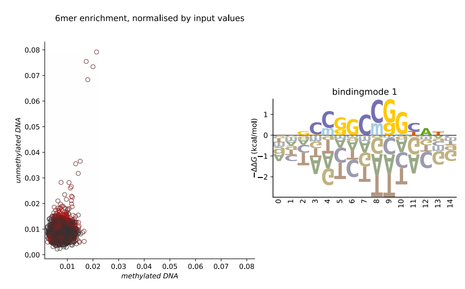


##### ZNF445 FL: methyl plus

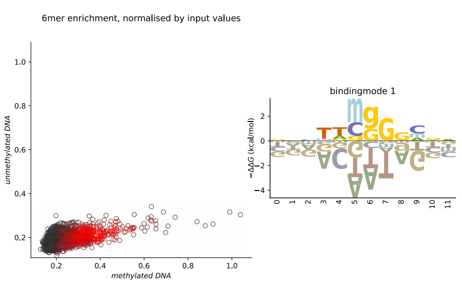


##### ZNF468 FL: no CG

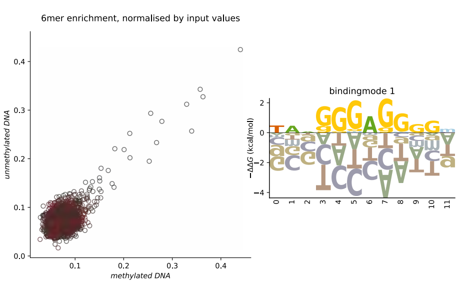


##### ZNF503 DBD: weak methyl plus

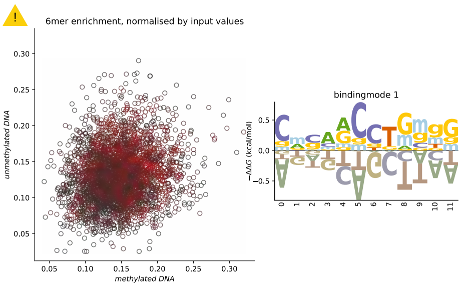


##### ZNF518B DBD: little effect/weak methyl plus

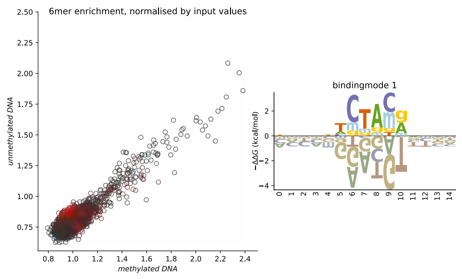


##### ZNF614 FL: weak methyl minus

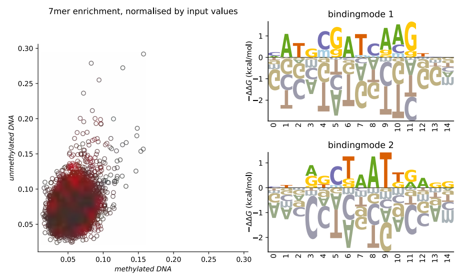


##### ZNF648 DBD: weak methyl plus

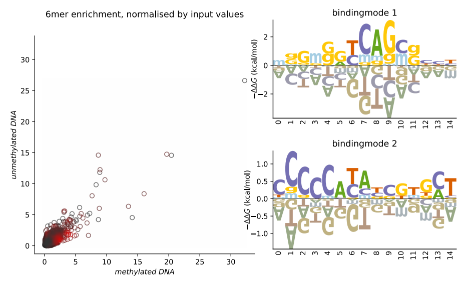


##### ZNF680 FL: little effect

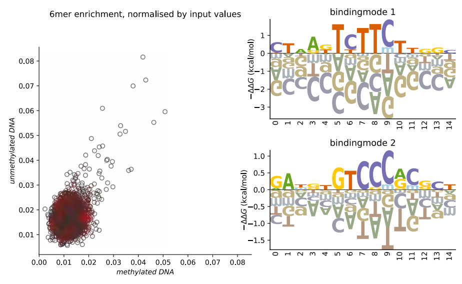


##### ZNF695 FL: little effect

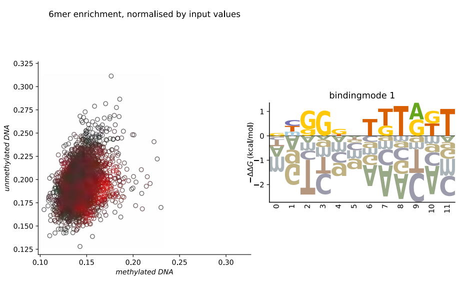


##### ZNF716 FL: methyl plus

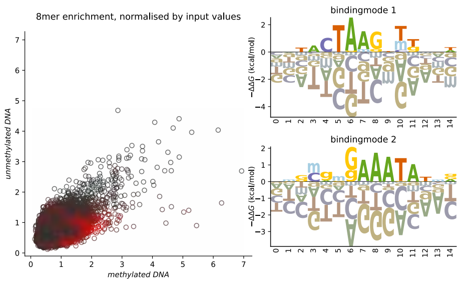


##### ZNF777 FL: methyl minus

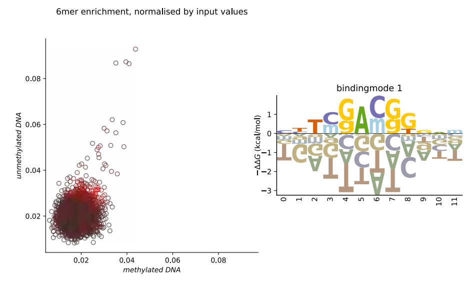


##### ZNF793 FL: little effect

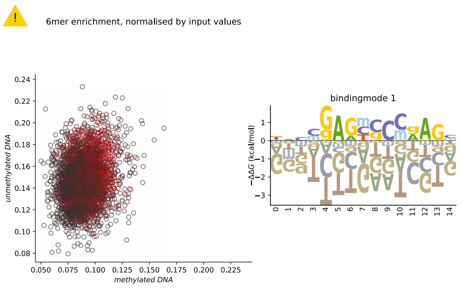

### C2H2 ZNF

##### ZNF831 DBD: little effect

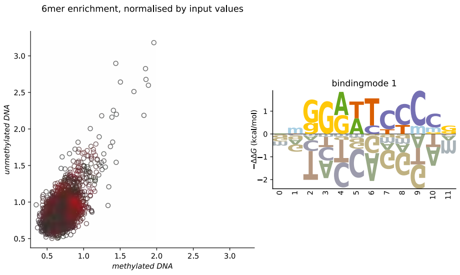


##### ZNF891 FL: little effect

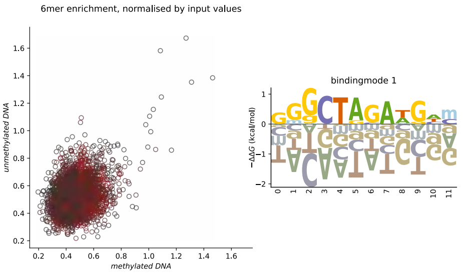


##### ZXDB DBD: weak methyl minus

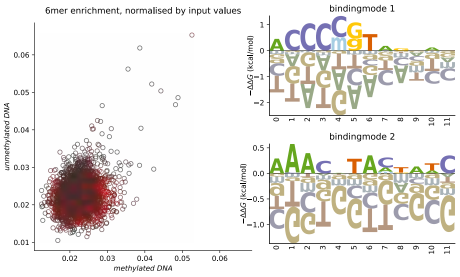


### Homeodomain, C2H2 ZNF

##### ZHX2 FL: methyl plus

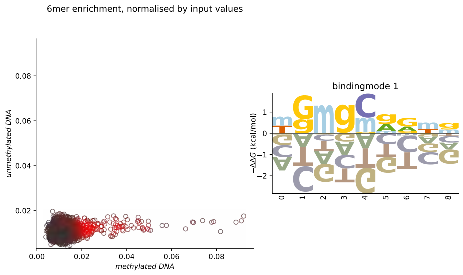


### BED ZF

##### FAM200B DBD: methyl minus

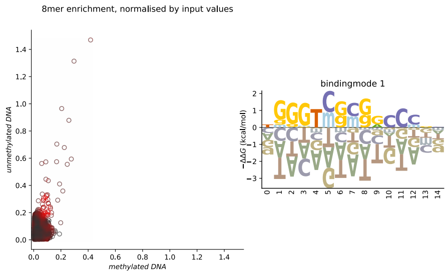


##### ZBED4 DBD: weak methyl minus

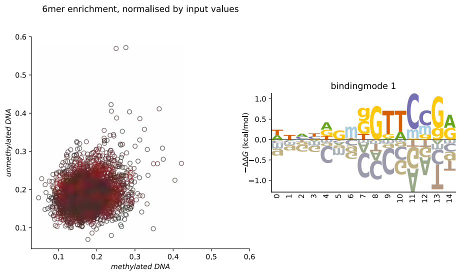


### unknown

##### TET2 FL: methyl plus

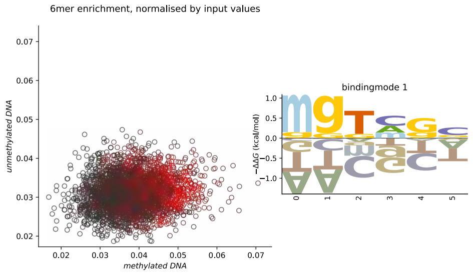

### bHLH

##### USF3 DBD: methyl minus

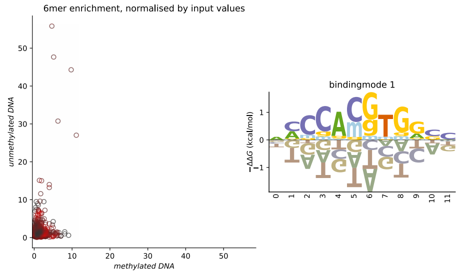


### CENPB

##### TIGD4 FL: weak methyl minus

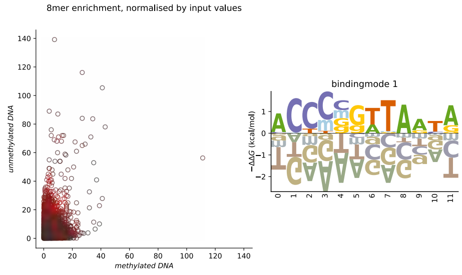


##### TIGD5 DBD:  methyl minus

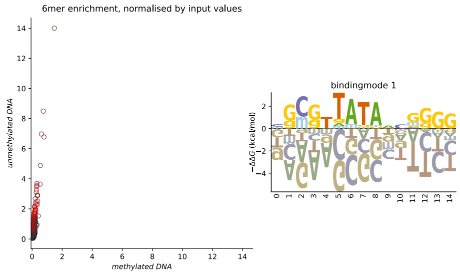


# CG-1

##### CAMTA1 DBD: little effect

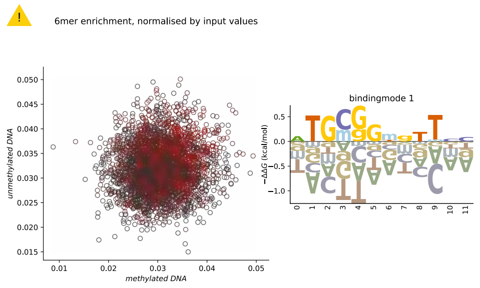


### CXXC ZF/MBD

##### CXXC4 FL: methyl minus

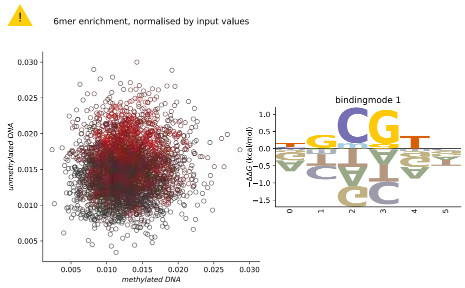


##### MBD1 DBD: methyl plus

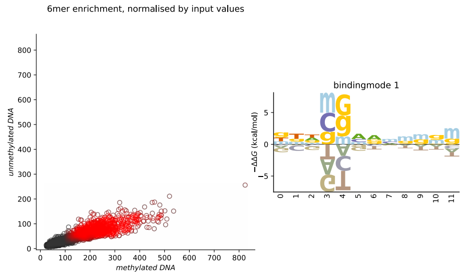


### MADF

##### MSANTD1 FL: no CG

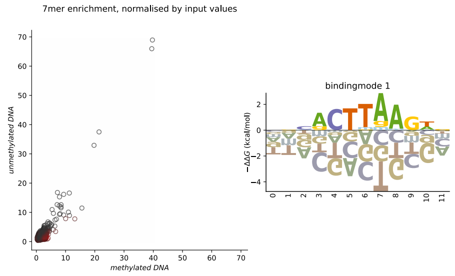


### Homeodomain/POU

##### TPRX1 DBD: no CG

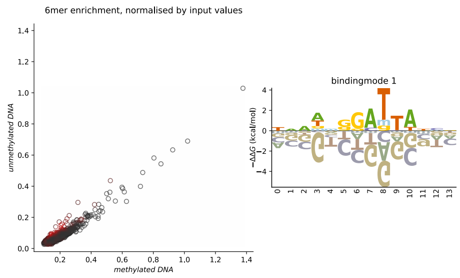


##### POU5F1 FL: methyl plus

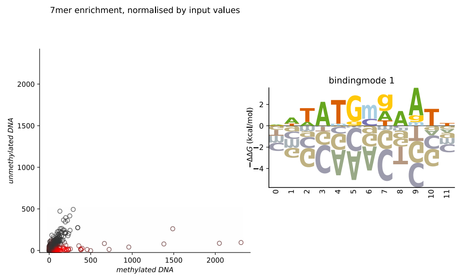


### Myb/SANT

##### TTF1 DBD: little effect

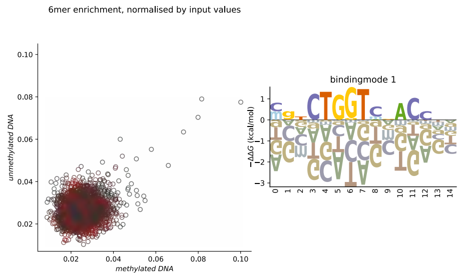


### RFX

##### RFX5 FL: methyl plus

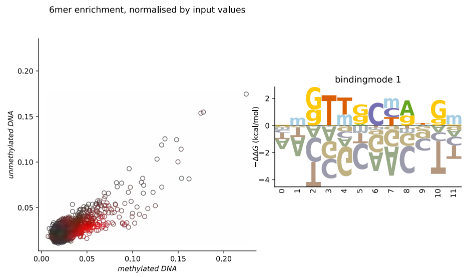
