## Supplementary Database 2 for "Identification of methylation-sensitive human transcription factors using meSMiLE-seq"

### Database 2.1

### Distribution of CG methylation levels of meSMiLE-seq inferred motifs in HEK293 cells.

ECDFs display the distribution of CG methylation levels within TF-specific motifs found in ChIP peaks, compared to the CG methylation distribution across the entire TF-specific ChIP peak.

Confidence intervals were estimated through bootstrapping, resampling 1,000 data points with replacement across 1,000 iterations.

WGBS data from three biological replicates were used, displayed are the median and the min/max confidence intervals.

*ChIP-seq data acquired from Razavi et al., DOI:10.1101/2024.11.11.622123*

*WGBS data for HEK293 acquired from Zhao S, Lu J, Pan B, Fan H et al. TNRC18 engages H3K9me3 to mediate silencing of endogenous retrotransposons. Nature 2023, GEO reference GSE233830.*

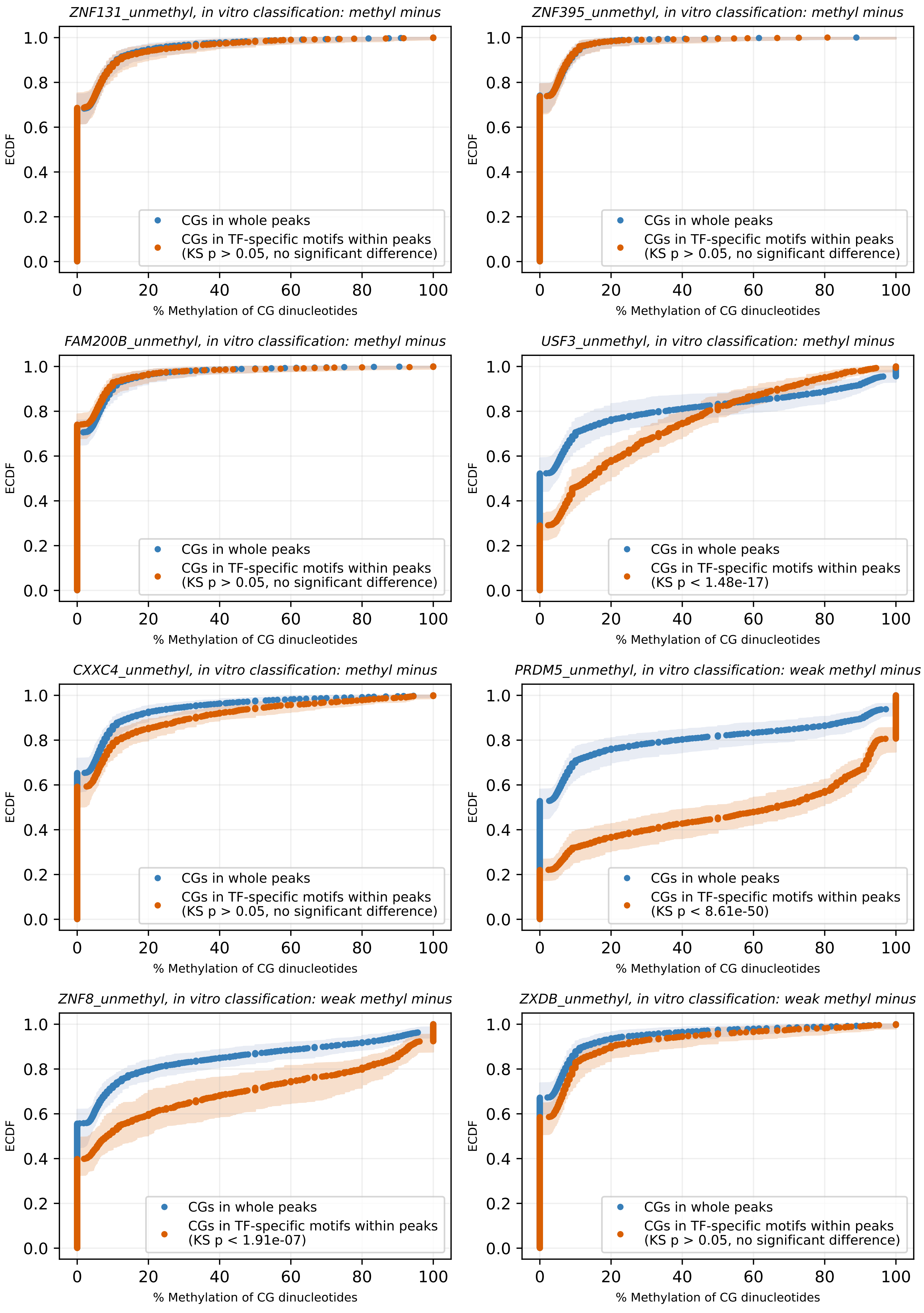

ZNF131_unmethyl, in vitro classification: methyl minus
CGs in whole peaks
CGs in TF-specific motifs within peaks
(KS p > 0.05, no significant difference)
ZNF395_unmethyl, in vitro classification: methyl minus
CGs in whole peaks
CGs in TF-specific motifs within peaks
(KS p > 0.05, no significant difference)
FAM200B_unmethyl, in vitro classification: methyl minus
CGs in whole peaks
CGs in TF-specific motifs within peaks
(KS p > 0.05, no significant difference)
USF3_unmethyl, in vitro classification: methyl minus
CGs in whole peaks
CGs in TF-specific motifs within peaks
(KS p < 1.48e-17)
CXXC4_unmethyl, in vitro classification: methyl minus
CGs in whole peaks
CGs in TF-specific motifs within peaks
(KS p > 0.05, no significant difference)
PRDM5_unmethyl, in vitro classification: weak methyl minus
CGs in whole peaks
CGs in TF-specific motifs within peaks
(KS p < 8.61e-50)
ZNF8_unmethyl, in vitro classification: weak methyl minus
CGs in whole peaks
CGs in TF-specific motifs within peaks
(KS p < 1.91e-07)
ZXDB_unmethyl, in vitro classification: weak methyl minus
CGs in whole peaks
CGs in TF-specific motifs within peaks
(KS p > 0.05, no significant difference)
ECDF
% Methylation of CG dinucleotides

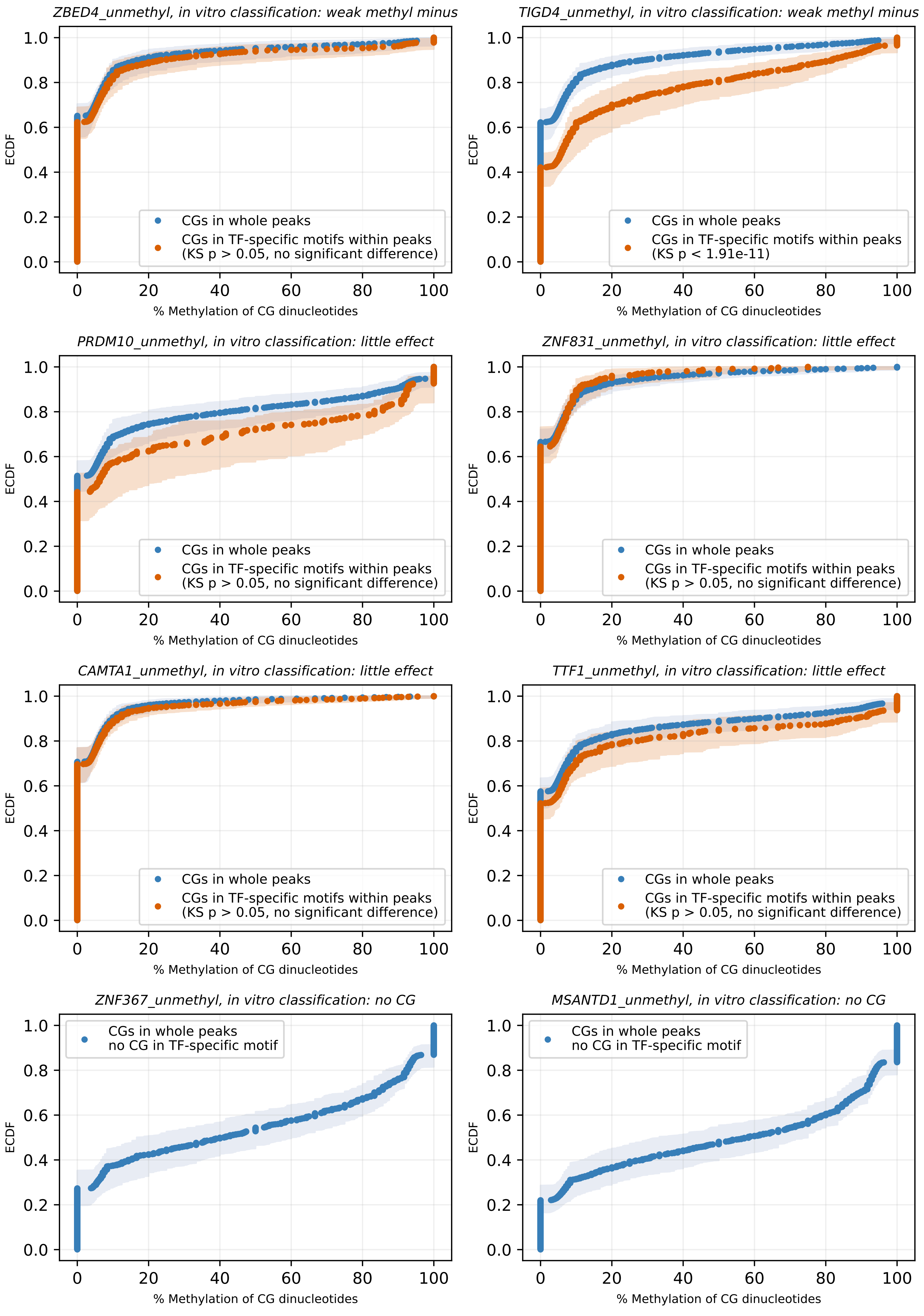
ZBED4_unmethyl, in vitro classification: weak methyl minus
ECDF
% Methylation of CG dinucleotides
CGs in whole peaks
CGs in TF-specific motifs within peaks
(KS p > 0.05, no significant difference)
TIGD4_unmethyl, in vitro classification: weak methyl minus
ECDF
% Methylation of CG dinucleotides
CGs in whole peaks
CGs in TF-specific motifs within peaks
(KS p < 1.91e-11)
PRDM10_unmethyl, in vitro classification: little effect
ECDF
% Methylation of CG dinucleotides
CGs in whole peaks
CGs in TF-specific motifs within peaks
(KS p > 0.05, no significant difference)
ZNF831_unmethyl, in vitro classification: little effect
ECDF
% Methylation of CG dinucleotides
CGs in whole peaks
CGs in TF-specific motifs within peaks
(KS p > 0.05, no significant difference)
CAMTA1_unmethyl, in vitro classification: little effect
ECDF
% Methylation of CG dinucleotides
CGs in whole peaks
CGs in TF-specific motifs within peaks
(KS p > 0.05, no significant difference)
TTF1_unmethyl, in vitro classification: little effect
ECDF
% Methylation of CG dinucleotides
CGs in whole peaks
CGs in TF-specific motifs within peaks
(KS p > 0.05, no significant difference)
ZNF367_unmethyl, in vitro classification: no CG
ECDF
% Methylation of CG dinucleotides
CGs in whole peaks
no CG in TF-specific motif
MSANTD1_unmethyl, in vitro classification: no CG
ECDF
% Methylation of CG dinucleotides
CGs in whole peaks
no CG in TF-specific motif

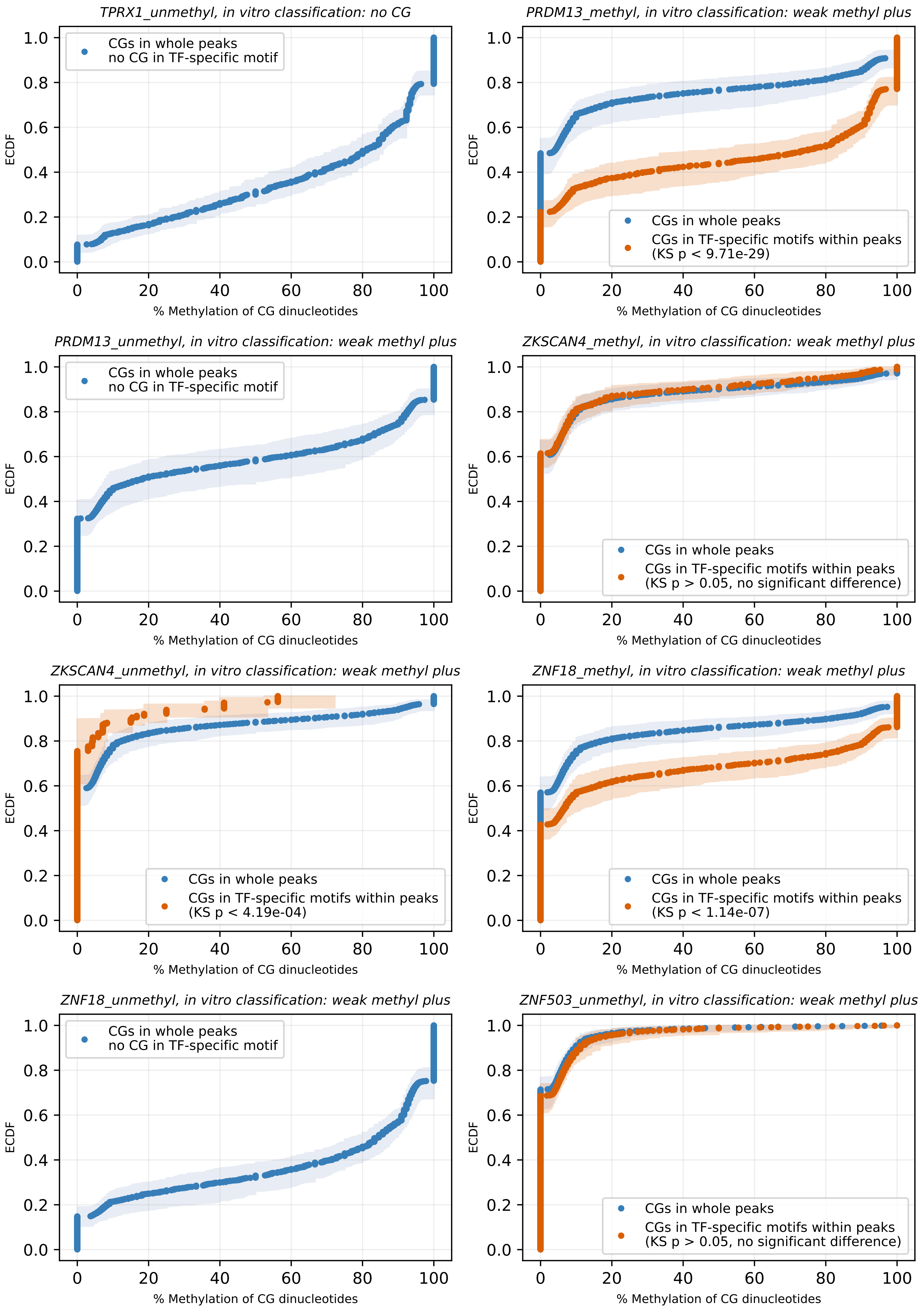

TPRX1_unmethyl, in vitro classification: no CG
CGs in whole peaks
no CG in TF-specific motif
PRDM13_methyl, in vitro classification: weak methyl plus
CGs in whole peaks
CGs in TF-specific motifs within peaks
(KS p < 9.71e-29)
PRDM13_unmethyl, in vitro classification: weak methyl plus
CGs in whole peaks
no CG in TF-specific motif
ZKSCAN4_methyl, in vitro classification: weak methyl plus
CGs in whole peaks
CGs in TF-specific motifs within peaks
(KS p > 0.05, no significant difference)
ZKSCAN4_unmethyl, in vitro classification: weak methyl plus
CGs in whole peaks
CGs in TF-specific motifs within peaks
(KS p < 4.19e-04)
ZNF18_methyl, in vitro classification: weak methyl plus
CGs in whole peaks
CGs in TF-specific motifs within peaks
(KS p < 1.14e-07)
ZNF18_unmethyl, in vitro classification: weak methyl plus
CGs in whole peaks
no CG in TF-specific motif
ZNF503_unmethyl, in vitro classification: weak methyl plus
CGs in whole peaks
CGs in TF-specific motifs within peaks
(KS p > 0.05, no significant difference)
ECDF
% Methylation of CG dinucleotides

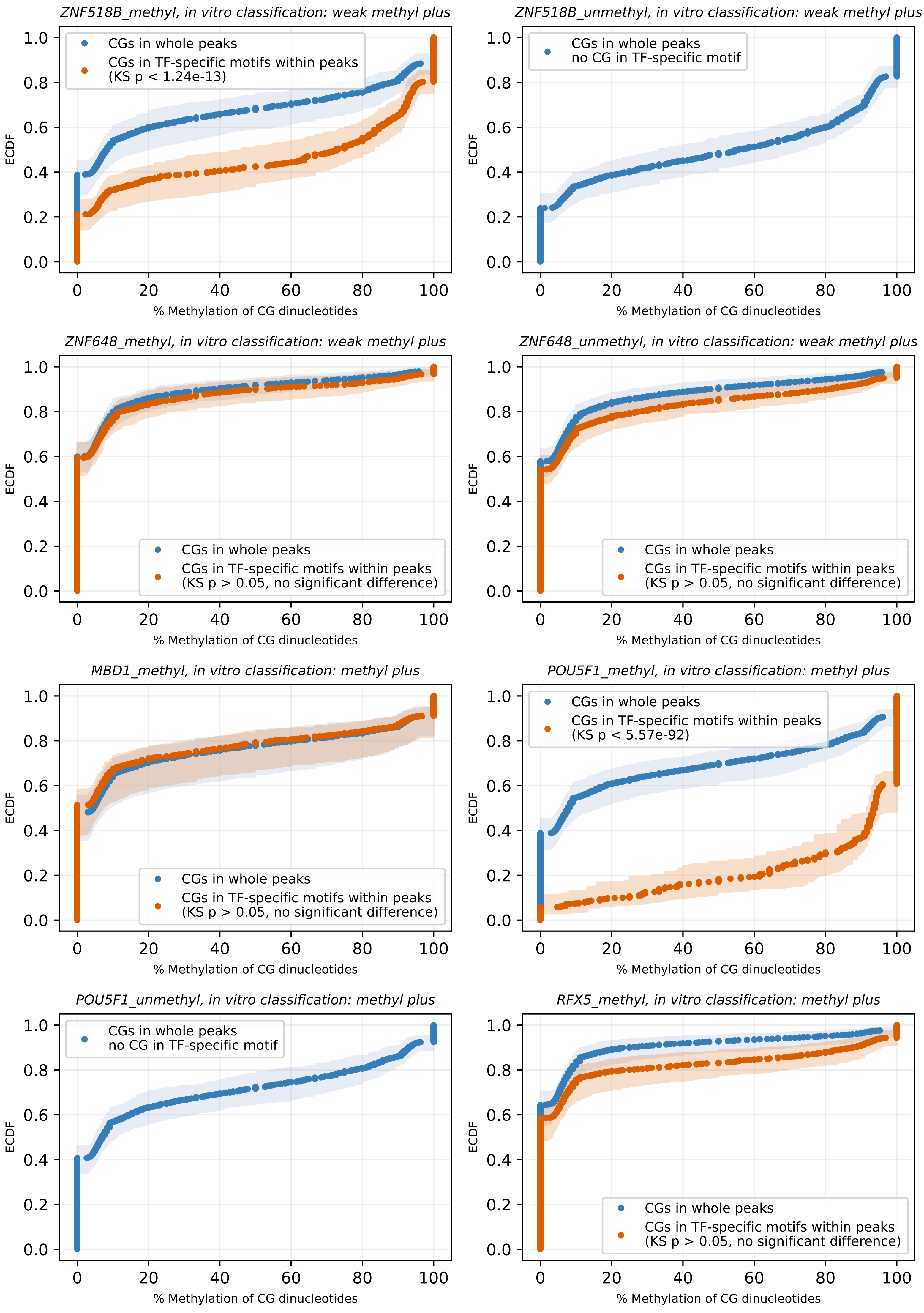

ZNF518B_methyl, in vitro classification: weak methyl plus
CGs in whole peaks
CGs in TF-specific motifs within peaks
(KS p < 1.24e-13)
ECDF
% Methylation of CG dinucleotides
ZNF518B_unmethyl, in vitro classification: weak methyl plus
CGs in whole peaks
no CG in TF-specific motif
ZNF648_methyl, in vitro classification: weak methyl plus
CGs in whole peaks
CGs in TF-specific motifs within peaks
(KS p > 0.05, no significant difference)
ZNF648_unmethyl, in vitro classification: weak methyl plus
MBD1_methyl, in vitro classification: methyl plus
POU5F1_methyl, in vitro classification: methyl plus
(KS p < 5.57e-92)
POU5F1_unmethyl, in vitro classification: methyl plus
RFX5_methyl, in vitro classification: methyl plus

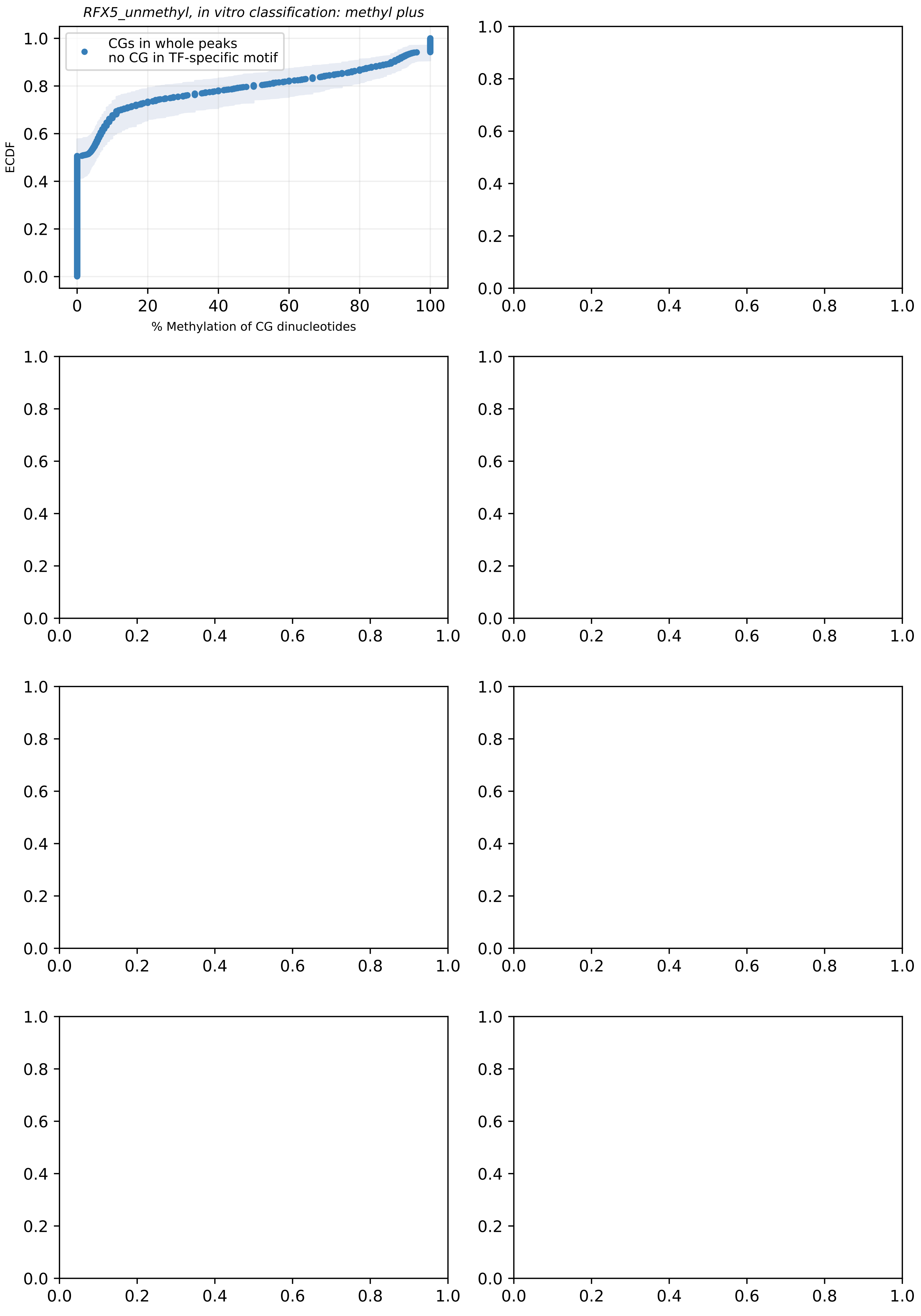
RFX5_unmethyl, in vitro classification: methyl plus
CGs in whole peaks
no CG in TF-specific motif
ECDF
% Methylation of CG dinucleotides

### Database 2.2

### Distribution of CG methylation levels of meSMiLE-seq inferred motifs in HEK293T cells.

ECDFs display the distribution of CG methylation levels within TF-specific motifs found in ChIP peaks, compared to the CG methylation distribution across the entire TF-specific ChIP peak.

Confidence intervals were estimated through bootstrapping, resampling 1,000 data points with replacement across 1,000 iterations.

WGBS data from four biological replicates were used, displayed are the median and the min/max confidence intervals.

*ChIP-exo data acquired from Imbeault M, Helleboid PY, Trono D.*
*KRAB zinc-finger proteins contribute to the evolution of gene regulatory networks.*
*Nature 2017, GEO reference GSE78099.*

*WGBS data for HEK293T acquired from Nuñez JK, Chen J, Pommier GC, Cogan JZ et al.*
*Genome-wide programmable transcriptional memory by CRISPR-based epigenome editing.*
*Cell 2021, GEO reference GSE168012.*

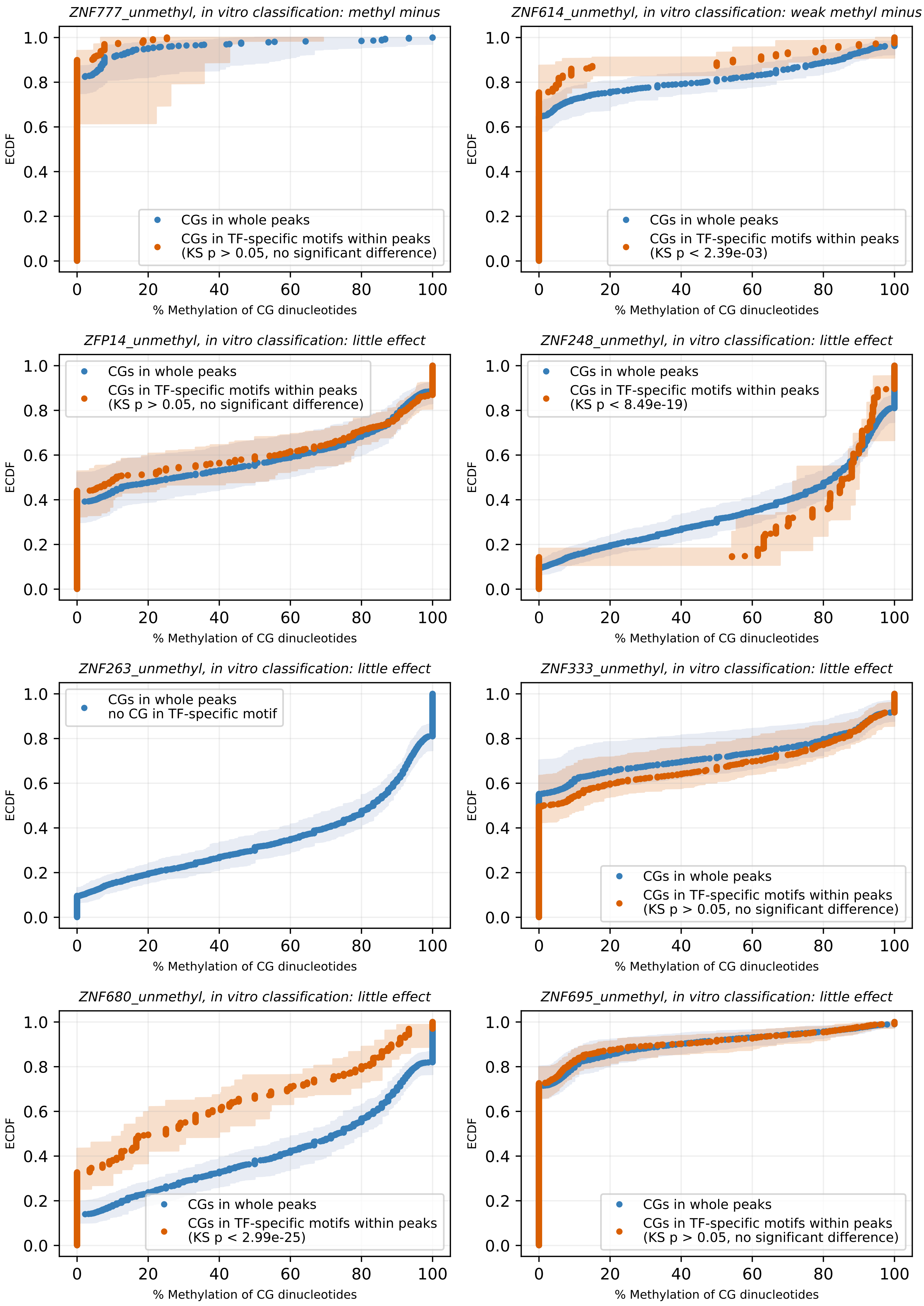
ZNF777_unmethyl, in vitro classification: methyl minus
CGs in whole peaks
CGs in TF-specific motifs within peaks
(KS p > 0.05, no significant difference)
ZNF614_unmethyl, in vitro classification: weak methyl minus
CGs in whole peaks
CGs in TF-specific motifs within peaks
(KS p < 2.39e-03)
ZFP14_unmethyl, in vitro classification: little effect
CGs in whole peaks
CGs in TF-specific motifs within peaks
(KS p > 0.05, no significant difference)
ZNF248_unmethyl, in vitro classification: little effect
CGs in whole peaks
CGs in TF-specific motifs within peaks
(KS p < 8.49e-19)
ZNF263_unmethyl, in vitro classification: little effect
CGs in whole peaks
no CG in TF-specific motif
ZNF333_unmethyl, in vitro classification: little effect
CGs in whole peaks
CGs in TF-specific motifs within peaks
(KS p > 0.05, no significant difference)
ZNF680_unmethyl, in vitro classification: little effect
CGs in whole peaks
CGs in TF-specific motifs within peaks
(KS p < 2.99e-25)
ZNF695_unmethyl, in vitro classification: little effect
CGs in whole peaks
CGs in TF-specific motifs within peaks
(KS p > 0.05, no significant difference)
ECDF
% Methylation of CG dinucleotides

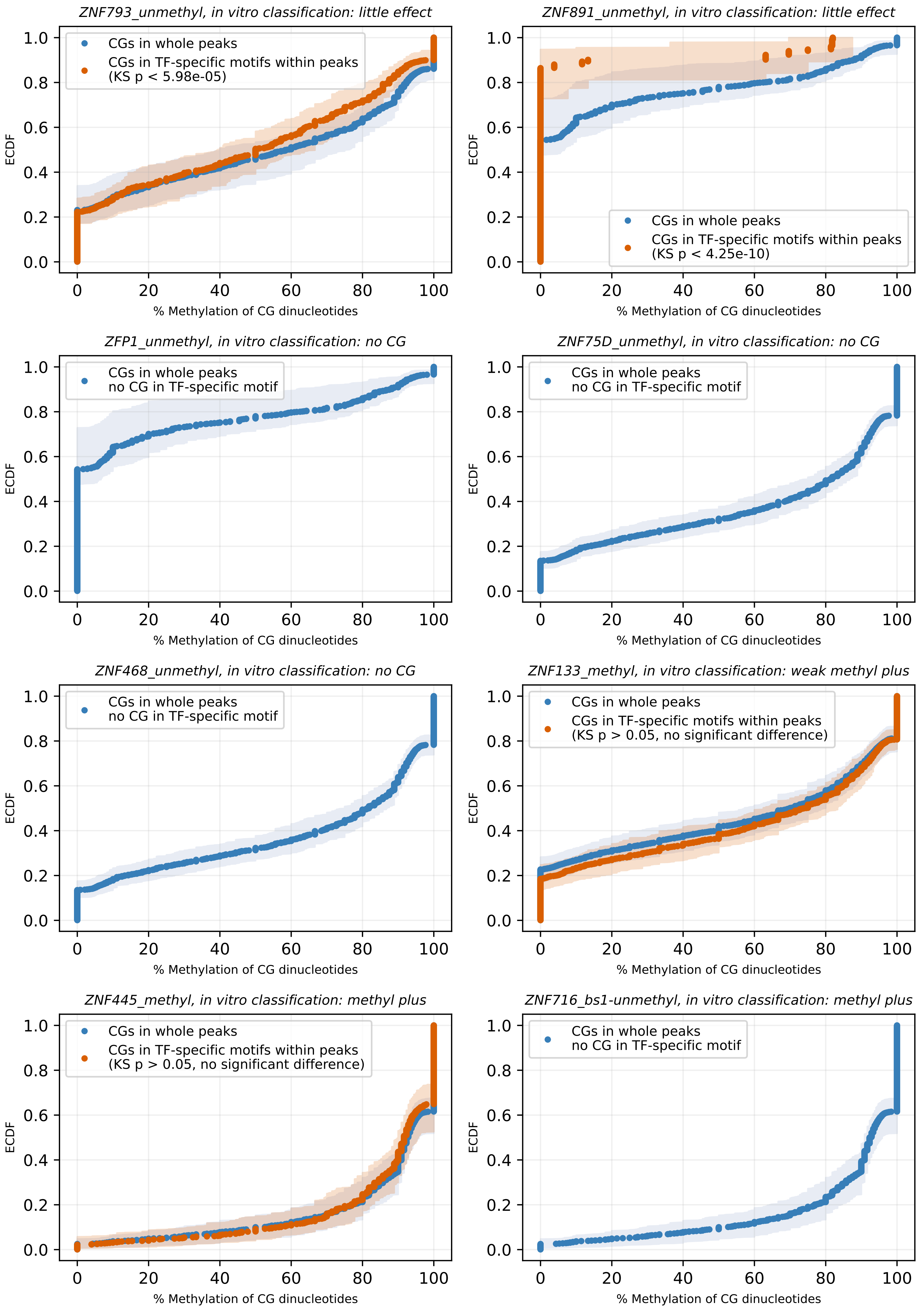
ZNF793_unmethyl, in vitro classification: little effect
CGs in whole peaks
CGs in TF-specific motifs within peaks
(KS p < 5.98e-05)
ZNF891_unmethyl, in vitro classification: little effect
CGs in whole peaks
CGs in TF-specific motifs within peaks
(KS p < 4.25e-10)
ZFP1_unmethyl, in vitro classification: no CG
CGs in whole peaks
no CG in TF-specific motif
ZNF75D_unmethyl, in vitro classification: no CG
CGs in whole peaks
no CG in TF-specific motif
ZNF468_unmethyl, in vitro classification: no CG
CGs in whole peaks
no CG in TF-specific motif
ZNF133_methyl, in vitro classification: weak methyl plus
CGs in whole peaks
CGs in TF-specific motifs within peaks
(KS p > 0.05, no significant difference)
ZNF445_methyl, in vitro classification: methyl plus
CGs in whole peaks
CGs in TF-specific motifs within peaks
(KS p > 0.05, no significant difference)
ZNF716_bs1-unmethyl, in vitro classification: methyl plus
CGs in whole peaks
no CG in TF-specific motif
ECDF
% Methylation of CG dinucleotides

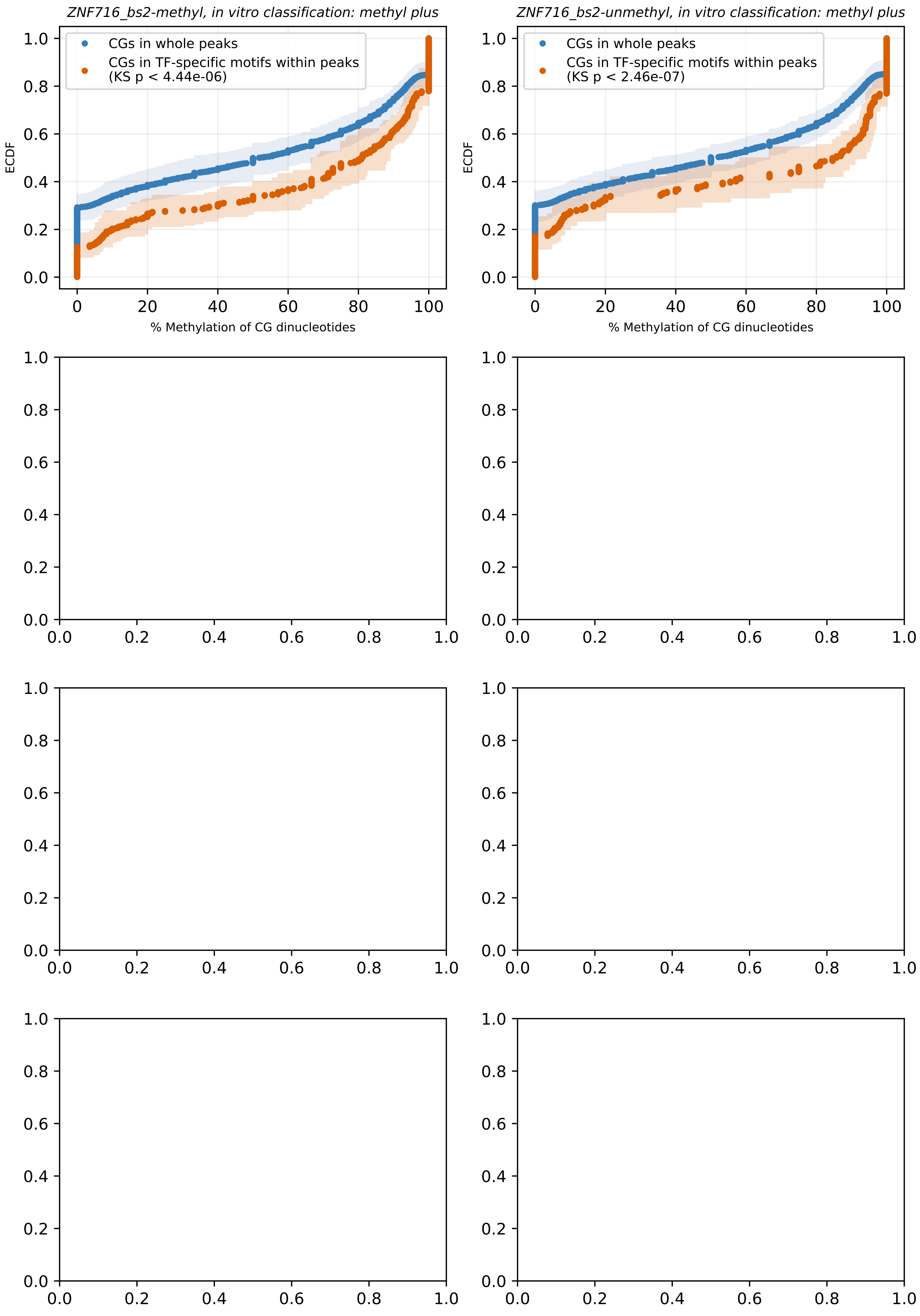
ZNF716_bs2-methyl, in vitro classification: methyl plus
CGs in whole peaks
CGs in TF-specific motifs within peaks
(KS p < 4.44e-06)
ECDF
% Methylation of CG dinucleotides
ZNF716_bs2-unmethyl, in vitro classification: methyl plus
CGs in whole peaks
CGs in TF-specific motifs within peaks
(KS p < 2.46e-07)
ECDF
% Methylation of CG dinucleotides
